# Molecular features of Myosin F adapted for driving actin flows in *Toxoplasma gondii*

**DOI:** 10.1101/2025.10.02.679852

**Authors:** Thomas E. Sladewski, Aoife T. Heaslip

## Abstract

*Toxoplasma gondii* (*T. gondii*) is a single-celled Apicomplexan parasite that relies on a highly polarized endomembrane system for its invasion into and survival within host cells. Recent advancements in imaging technologies have revealed that vesicle transport and organization of organelles in the endomembrane pathway requires a highly dynamic actin cytoskeleton. These dynamics in turn rely on the activity of Myosin F (MyoF), a molecular motor unique to Alveolates. The defining characteristic of this molecular motor is a WD40 beta-propeller domain, exclusively found in this class of myosin. To understand the mechanism by which MyoF controls the dynamics and organization of actin, we studied the biophysical properties of the purified motor *in vitro*. A MyoF construct lacking its WD40 tail domain (MyoFΔtail) is dimeric and can bind and translocate actin in an *in vitro* motility assay. Single molecule studies show that the dimeric construct is non-processive however small ensembles move inefficiently on single filaments of skeletal actin. In contrast, single molecules of the full-length motor move processively on *Toxoplasma* actin and jasplakinolide-stabilized skeletal actin bundles. Electron microscopy of negatively stained images of MyoF and quantitative size exclusion chromatography shows that the WD40 domain oligomerizes to form a complex containing multiple dimeric molecules, which provides an explanation for why the full-length motor is processive compared to the dimeric MyoFΔtail construct. Finally, we show that MyoF binds microtubules through its WD40 domain and can slide actin filaments relative to microtubules. We propose a model whereby MyoF oligomers drive actin dynamics by translocating filaments relative to the parasite’s cytoskeleton. These molecule features provide new insight into how MyoF functions in the cell to regulate actin organization during vesicle transport.

## Introduction

*Toxoplasma gondii* is an obligate intracellular parasite that infects approximately one-third of the world’s population. Infection results in severe disease in immunocompromised individuals and when infection occurs *in utero* (1, 2). Disease pathogenesis and parasite survival depend on the parasite’s ability to complete multiple rounds of its lytic cycle. This involves host cell invasion, replication, and egress, which ultimately destroys the invaded host cell (3). These steps rely on a highly polarized secretion system made up of micronemes, rhoptries which are localized at the apical end of the parasite, and dense granules which are highly motile vesicles distributed throughout the cytosol (4). The secreted contents of the three vesicle types allows the parasite to attach and invade host cells (5, 6), modulate host immune response pathways (7), establish a chronic infection (8), and permeabilize cell membranes to facilitate egress (9).

We previously determined that the organization of the endomembrane system in *T. gondii* is controlled by a divergent actin gene (*Tg*Act1) and myosin F (MyoF), a class of myosin motor found exclusively in alveolate protists, which encompasses the Apicomplexan phylum (10). Depletion of either of these proteins results in defects in the movement of dense granules, Rab6, Rab11a, and Rop1-positive vesicles (4, 11, 12). Other components of the endomembrane system are also affected including Golgi morphology and positioning, inheritance of a plastid-like organelle called the apicoplast, and ER tubule movement (4, 12, 13).

It is our goal to determine the mechanisms by which *Tg*Act1 and MyoF control the positioning of such a wide range of cellular cargoes in *T. gondii*. Live cell imaging using an actin chromobody revealed that intracellular parasites contain a highly dynamic cytosolic actin network (14-16). Depletion of MyoF results in severely reduced actin dynamics and compaction of the cytosolic actin network into bundled structures apical to the Golgi (16). The requirement of MyoF for filament dynamics indicates that this motor and *Tg*Act1 make up an unconventional actomyosin system that drives cargo transport in *T. gondii* which is distinct from the “canonical” cargo transport described in yeast and mammals whereby molecular motors bind cargo via the C-terminal tail domains and transport it along actin or microtubule cytoskeletal tracks (17-20). In this study, we analyze the molecular features and biophysical properties of MyoF to understand how the motor facilitates actin dynamics and drives cargo transport.

## Results

### The speed of MyoF is dependent on light chain composition

MyoF has a well conserved motor domain with the exception of three unique inserts (**Fig. 1A-B**), the function of which are currently unknown (4). An alpha-fold model of the MyoF motor domain (blue) and lever arm bound to a single calmodulin light chain (grey) shows that the second unique insert in the motor domain is located adjacent to the light chain bound to the first IQ motif (**Fig. 1B**). The lever arm region of MyoF contains six putative light-chain binding IQ motifs (**Fig. 1C**). The dimerization domain of MyoF is predicted to begin within the last IQ motif and contains a total of 22 heptads with the potential to form an α-helical coiled-coil spanning amino acids I974 to L1174 (**Fig. 1D**). While the ability of an alpha helix to form a coiled-coil is dependent on the amino acid composition within the heptad repeat, 22 heptads should, in principle, form a stable dimer, given that the first 20 heptads of vertebrate myosin Va are sufficient for dimerization (18). An alpha fold model of the coiled-coil region (E993 to R1204) predicts an additional 77 amino acids of coiled-coil sequence, with a break in the middle of the rod region (**Fig. 1E**). This may prevent the rod from forming an extended conformation, typical of some cargo transporting molecular motors such as myosin Va or kinesin-1 (17, 20). The coiled-coil region is followed by ∼400 amino acids of unknown structure, terminating with a seven-bladed WD40 beta-propeller domain (**Fig. 1F**), the defining feature of this class of myosin motor.

**Figure 1:**
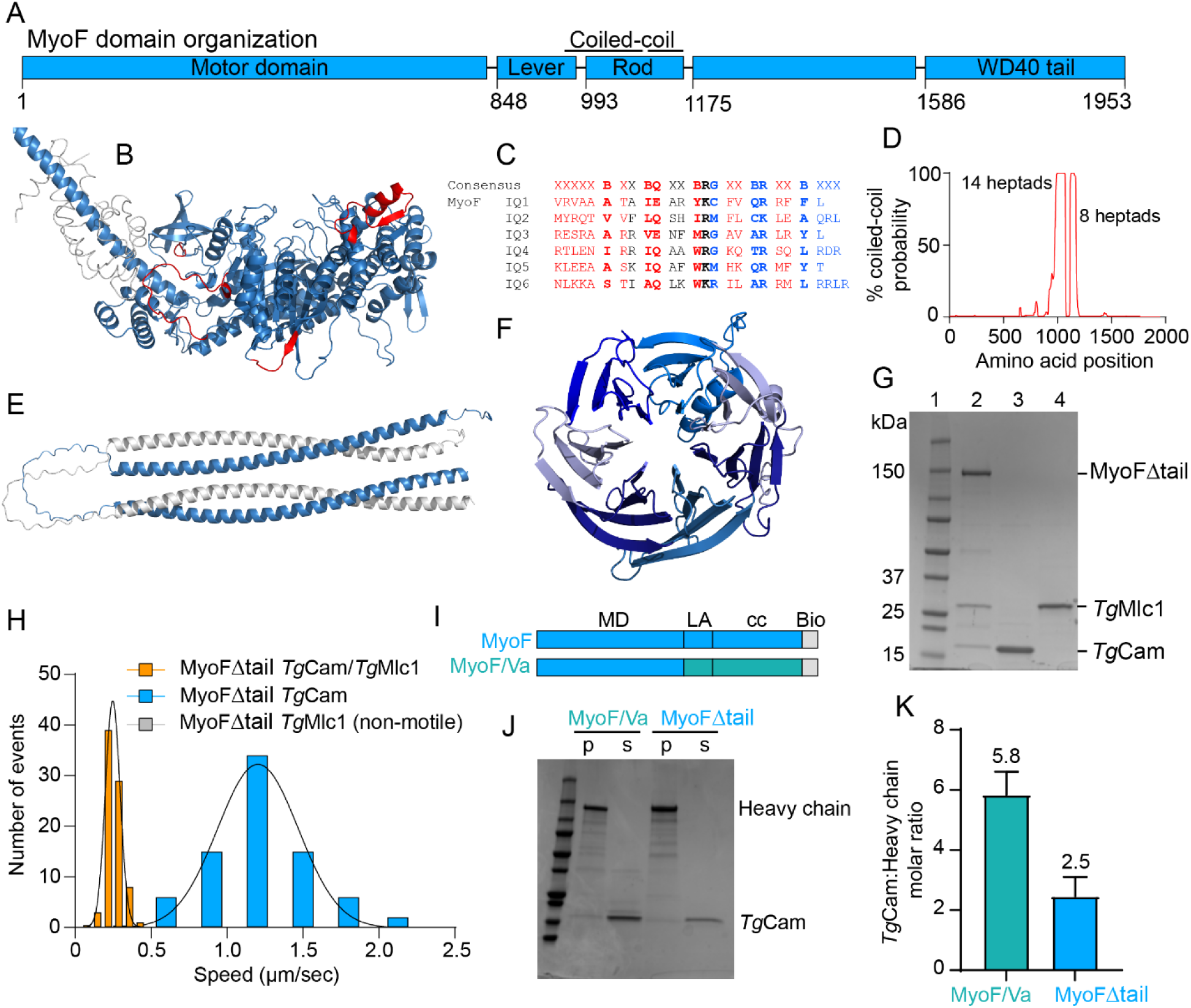
Analysis of MyoF domain organization and light chain binding. (**A**) Domain organization of MyoF. Region predicted to form a coiled-coil is indicated. Amino acids 1175-1585 are sequence with unknown structure. (**B**) Alpha fold model of the MyoF motor domain (a.a. 1-847) and a portion of the lever arm (a.a. 848-885) (blue) showing four unique inserts (red). Calmodulin (grey) (PDB 2DFS) is modeled onto the first IQ motif sequence. (**C**) Aligned sequences of the six predicted IQ motifs (consensus sequence IQxxxRGxxxR) in the MyoF lever arm (x, any residue; B, hydrophobic; red, C-lobe interacting; blue, N-lobe interacting). (**D**) Coiled-coil probability (Ncoils) of the MyoF sequence. Number of heptads are calculated from amino acid sequences with a p-value greater than 0.5 (50%) (I974 – L1174). (**E**) Predicted structure of the coiled-coil rod region of MyoF (E993 – W1251) using AlphaFold. (**F**) Structure of the WD40 tail domain (a.a. M1572-A1941) predicted by alpha fold. Unstructured C-terminal amino acids G1942-V1952 are not shown. (**G**) Coomassie-stained SDS-PAGE gel of (lane 1) protein molecular weight marker; (lane 2) FLAG affinity purified MyoFΔtail from *Sf9* cells co-expressed with *Tg*Cam and *Tg*Mlc1; (lane 3) HIS purified *Tg*Cam and (lane 4) *Tg*Mlc1 expressed from *E. coli*. (**H**) Gliding filament *in vitro* motility speed distributions of MyoFΔtail bound to *Tg*Cam and *Tg*Mlc1 (orange, mean ± SD, 0.24 ± 0.05 µm/sec, n=80) and MyoFΔtail bound to only *Tg*Cam (blue, mean ± SD, 1.2 ± 0.27 µm/sec, n=78). (**I**) Domain organization of MyoFΔtail and MyoF/Va constructs showing MyoF sequences in blue and mammalian myosin Va sequences in teal. (**J**) Coomassie-stained SDS-PAGE gel showing supernatant (s) and pellet (p) fractions of centrifuged heat-treated MyoFΔtail and MyoF/Va constructs. (**K**) Average calmodulin:heavy chain ratio for MyoF/Va (teal) and MyoFΔtail constructs (N=3).

Light chains function to mechanically stabilize the lever arm of myosin which is required for motility. Thus, to study the motile properties of recombinant MyoF *in vitro*, we first identified its native light chains by performing immunoprecipitation (IP) assays using GFP trap affinity resin and lysates from a parasite line where the motor was fused to a C-terminal EmeraldFP (EmFP) tag (4). A fraction of the eluted protein showed the presence of a ∼200 kDa band which likely corresponds to MyoF-EmFP, and bands approximating 60 and 15 kDa as shown by silver staining (**Fig. S1A**). MyoF interacting proteins were subsequently identified using liquid chromatography-mass spectrometry (LC-MS) (**Table S1**). In two of the three IPs, two light chains were immunoprecipitated with MyoF: Calmodulin (*Tg*Cam) (ToxoDB accession code TGME49_249240) and myosin light chain (*Tg*Mlc1) (ToxoDB accession code TGME49_257680) (21).

Having identified the native light chains, we next purified recombinant MyoF using the baculovirus/*Sf9* cell system. Because the tail domain of other classes of myosins interact with the motor domain to form an auto-inhibited complex, we first purified a construct of MyoF lacking its WD40 tail domain (a.a. 1-1,265) which was fused to a C-terminal biotin-FLAG tag which is used to functionalize and affinity purification the protein (MyoFΔtail). The sequence of the biotin tag is derived from the biotin carboxyl carrier protein of acetyl-CoA carboxylase which becomes biotinylated *in vivo* (22, 23). This construct was co-expressed in *Sf9* cells with untagged *Tg*Cam and *Tg*Mlc1 light chains in a back-to-back system which ensures similar expression levels (**Fig. S1B**). All MyoF motor constructs in this study were also expressed back-to-back with a *T. gondii*-specific co-chaperone *Tg*UNC (ToxoDB accession code TGME49_249480) (**Fig. S1B**) (24, 25). While *Tg*UNC was not necessary for the expression of soluble MyoF, we found co-expression with the chaperone produced preparations with higher protein yields.

Using this expression strategy, we find that both *Tg*Cam and *Tg*Mlc1 co-purify with MyoFΔtail confirming their association with the motor (**Fig. 1G, lane 2, Fig. S2A**). To understand how *Tg*Cam and *Tg*Mlc1 differentially affect the activity of the MyoF, we purified the motor co-expressed with *Tg*Cam alone (**Fig. S2B**) and with *Tg*Mlc1 alone (**Fig. S2A**). Using an *in vitro* motility assay, we found that MyoFΔtail bound to *Tg*Cam and *Tg*Mlc1 glided filaments with an average speed of 0.24 µm/sec (**Fig. 1H, Movie S1**), while the gliding motility speed for MyoF bound to only *Tg*Cam was enhanced 5-fold (1.2 µm/sec) (**Fig. 1H, Movie S2**). MyoFΔtail bound to only *Tg*Mlc1 was non-motile (**Fig. 1H, Movie S3**). This data suggests that *Tg*Cam is sufficient to fully occupy and mechanically stiffen the lever arm of MyoF to support motility, and that *Tg*Mlc1 may serve a regulatory function. Sequence analysis shows that the lever arm region of MyoF has six putative light chain binding sites (**Fig. 1C**). To determine if all IQ motifs are capable of light chain binding, we counted the number of light chains using a calmodulin-binding assay described previously (26, 27). MyoFΔtail bound to *Tg*Cam was thermally denatured, transferred to ice, and then centrifuged at high speed to separate soluble and insoluble fractions. Because the heavy chain precipitates and *Tg*Cam remains soluble, the concentration of the supernatant and pellet can be used to calculate the number of *Tg*Cam light chins bound per heavy chain. As a control, we purified a chimeric construct from *Sf9* cells containing the MyoF motor domain and mammalian myosin Va lever arm and rod sequences (MyoF/Va), which is known to bind six calmodulin light chains (**Fig. 1I, Fig. S2D**). Using this strategy, we found that the chimeric MyoF/Va control binds approximately six light chains as expected (**Fig. 1J-K, teal**), while MyoFΔtail binds approximately three (**Fig. 1J-K, blue**) indicating that MyoF binds half as many light chains than myosin Va and fewer than predicted from the amino acid sequence.

### A single MyoF dimer is non-processive

A hallmark of yeast and mammalian cargo transporting myosins is the ability to move processively on actin filaments as a single molecule (20). To determine if a truncated dimer of MyoF is processive, we visualized the movement of MyoFΔtail on skeletal actin filaments bound to a quantum dot (Qdot). Streptavidin-coated Qdots were bound to MyoFΔtail through its C-terminal biotin tag at a ratio of 1 motor to 5 Qdots to ensure that the majority of Qdots are bound to a single motor. Under these mixing conditions, single molecules of MyoFΔtail bound to *Tg*Cam and *Tg*Mlc1 light chains did not move on actin (**Movie S4**). Because light chain composition has a dramatic effect on the speed of MyoFΔtail in the *in vitro* gliding filament motility assay, we tested whether removing *Tg*Mlc1 had an effect on the processivity of MyoFΔtail. We find that single molecules of MyoFΔtail bound to only *Tg*Cam also do not move on skeletal actin filaments (**Movie S5**). These results indicate that the single dimers of the truncated motor are unable to support motility and are therefore non-processive on skeletal actin.

We next tested if small motor ensembles of MyoFΔtail bound to a single Qdot can move continuously on actin filaments. MyoFΔtail was mixed with Qdots at a 10:1 molar ratio to ensure saturation of streptavidin binding sites. Given its geometry and occupancy, this allows a single Qdot to bind 4-6 myosin motors (28). We find that small teams of MyoFΔtail bound to *Tg*Cam and *Tg*Mlc1 are capable of supporting motility on skeletal actin filaments (**Movie S6**) with a characteristic run length (λ) of 0.65 µm (**Fig. 2A**) and average speed of 0.51 µm/sec (**Fig. 2B, orange**). To test whether removal of *Tg*Mlc1 changes the properties of small ensembles of truncated MyoF, we imaged the movement of multiple MyoFΔtail motors on skeletal actin bound to only *Tg*Cam. We find little difference in the run length (0.72 µm) of these ensembles compared to MyoF bound to both light chains (**Movie S7, Fig. 2C**). However, the average speed of movement was much faster (3.45 µm/sec), consistent with *in vitro* gliding filament motility results (**Fig. 1H**). Taken together, these data indicate that light chain composition affects the speed of MyoF but not the processivity or run length of the truncated MyoFΔtail construct.

**Figure 2:**
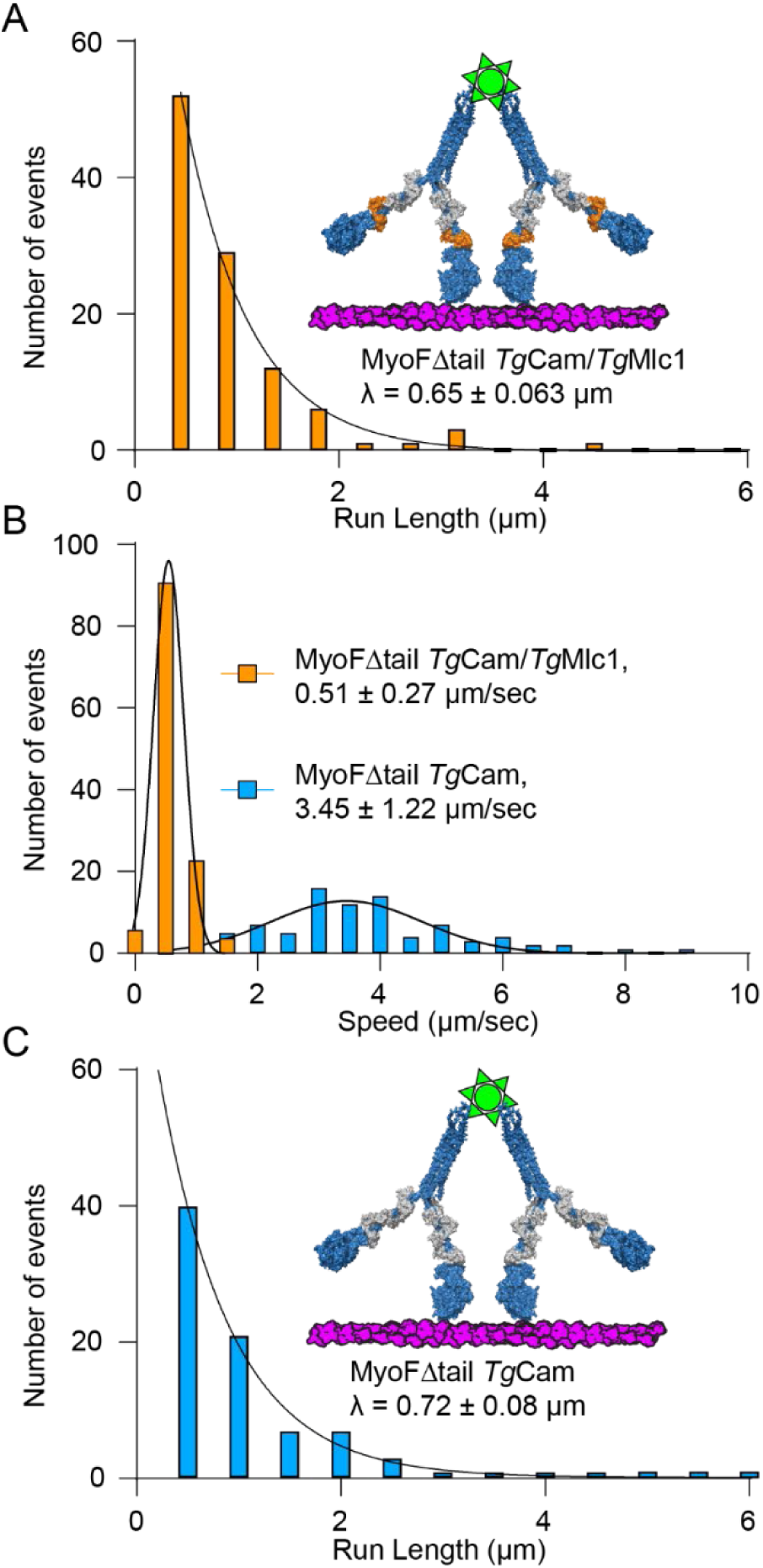
Effect of light chain composition on MyoF motility. (**A**) Frequency distribution of run lengths for multiple motors of MyoFΔtail co-purified with *Tg*Cam and *Tg*Mlc1 bound to a Qdot on skeletal actin (n = 124). Characteristic run length (λ) is shown ± SE of the fit. (**B**) Histograms comparing multiple motor run speeds MyoFΔtail co-purified with *Tg*Cam and *Tg*Mlc1 (orange) (n = 124) and MyoFΔtail co-purified with *Tg*Cam (blue) on skeletal actin (n = 84). Histograms were fit to a single Gaussian distribution to determine the average speed ± SD for each construct. (**C**) Frequency distribution of run lengths for multiple motors of MyoFΔtail co-purified with *Tg*Cam on skeletal actin (n = 124). Error is in SD.

### Teams of MyoFΔtail show optimal motility on actin-fascin bundles

Some classes of cargo transporting myosin motors with shortened lever arms are optimized for movement on actin bundles rather than single actin filaments (29-31). This property allows them to target bundled actin structures such as filopodium and stereocilia and become enriched at their tips (32-36). To determine if MyoF is targeted to actin bundles *in vivo*, we ectopically expressed the full-length motor in *Sf9* cells fused at its C-terminus with the mClover3 variant of GFP (MyoF-GFP) and imaged its localization using epifluorescence microscopy. We find that MyoF is strongly enriched to the ends of actin-enriched filopodium (**Fig. S3**).

Its ectopic localization and shortened lever arm suggest that MyoF may move better on parallel actin bundles which contain additional lateral binding sites that could enhance motility by ensuring at least one head remains bound to the actin. We compared the movement of teams of MyoFΔtail bound to a single Qdot on single actin filaments and actin-fascin which crosslinks actin filaments into parallel bundles that are separated by 9 nm (37). We find that teams of MyoFΔtail have a relatively low run frequency on single actin filaments (**Fig. 3A, left, Fig. 3B**, **Fig. 3C**). However, the frequency of movement and run length was dramatically enhanced on actin bundles (**Fig. A, right, Fig. C, Fig. D**). The average speed of movement on bundles is reduced (**Fig. 3D**) which may result from lateral movements and side stepping, a behavior seen with other molecular motors that are optimized to move on bundles (38). To test whether single molecules of MyoFΔtail move on bundles, we mixed clarified motors with Qdots at a 1:5 ratio to ensure only one motor is bound per Qdot and imaged its motility using TIRF microscopy on actin-fascin bundles. When we do this, no runs were observed (**Movie S8**) indicating that single molecules of MyoFΔtail are non-processive on actin bundles.

**Figure 3:**
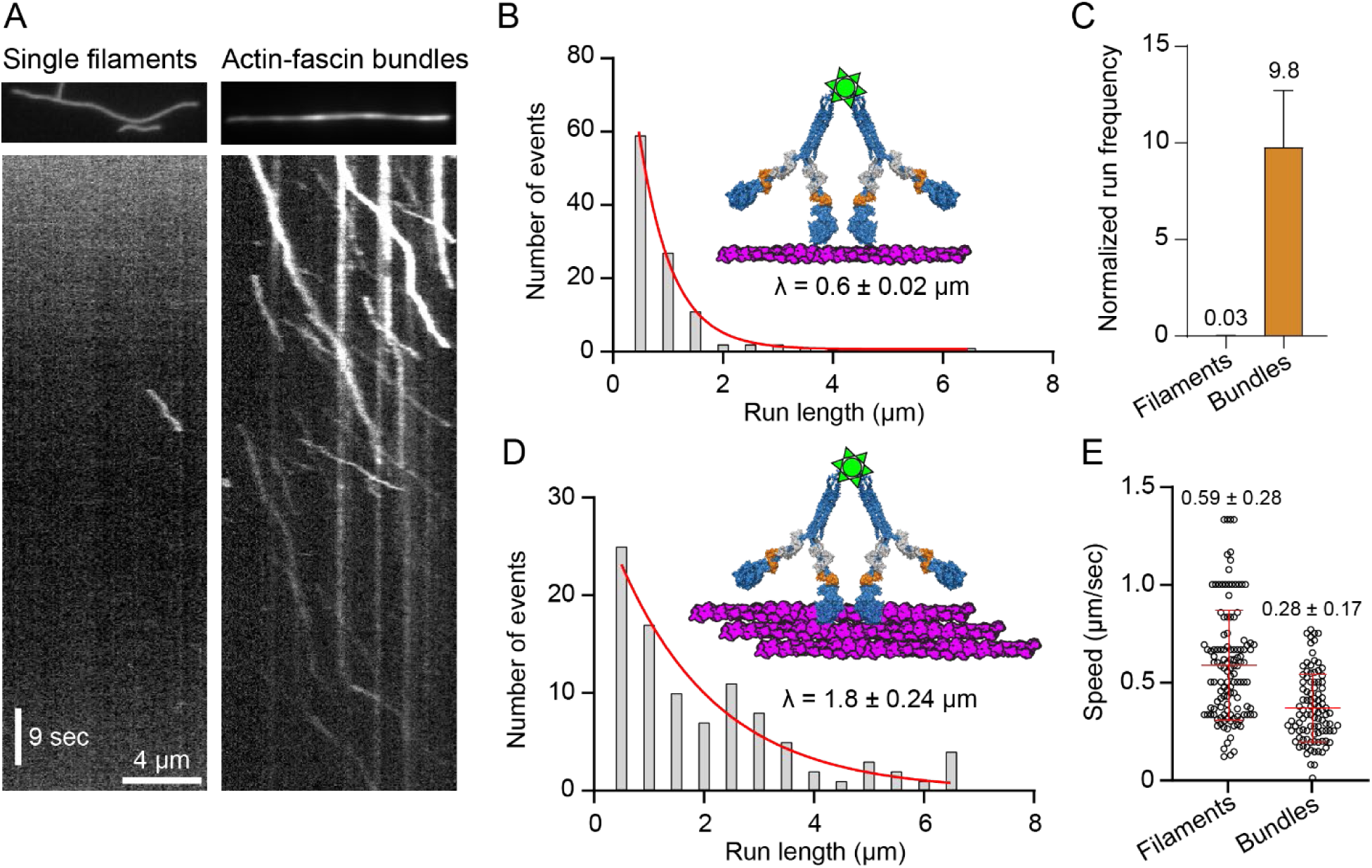
MyoFΔtail motility on single actin filaments and actin-fascin bundles. (**A**) Kymograph (distance vs. time) showing the movement (diagonal lines) of multiple motors of MyoFΔtail co-purified with *Tg*Cam and *Tg*Mlc1 on single skeletal actin filaments (left) vs. actinfascin bundles (right) at the same concentration of motor. (B) Frequency distribution of run lengths for multiple motors of MyoFΔtail on skeletal actin (n = 124). Characteristic run length (λ) is shown ± SE of the fit. (**C**) Comparison of the mean run frequency for multiple motors of MyoFΔtail on single skeletal actin filaments (mean ± SD, 0.03 ± 0.01, n = 22 filaments) versus MyoFΔtail on actin-fascin bundles (mean ± SD, 0.03 ± 0.01, n = 8 bundles). Event frequency was normalized per µM Qdot per µm actin track per s, as done previously (38). (**D**) Run length frequency distribution of multiple motors of MyoFΔtail on actin-fascin bundles (n = 99). The characteristic run length (λ) on bundles is significantly longer than on filaments, P < 0.001, using the Kolmogorov-Smirnov Test. (**E**) Run speed distributions comparing multiple motors of MyoFΔtail on single actin filaments versus actin-fascin bundles. Mean run speed ± SD is shown. Average run speeds are significantly different (P value < 0.0001, using a two-tailed t-test).

### The WD40 tail domain oligomerizes MyoF

We next used electron microscopy (EM) to determine the stoichiometry and organization of MyoF. Negatively stained EM images of the MyoFΔtail construct were compared to the MyoF/Va chimera because the structure of the MyoVa backbone is well established (18). MyoFΔtail shows “V-shaped” particles containing pairs of 10 nm diameter heads consistent with a dimer (**Fig. 4A, white arrowheads, blue**). Unlike MyoF/Va, which has an extended lever arm (**Fig. 4B, yellow arrowheads**), MyoF has a lever arm approximately half the length (**Fig. 4A, yellow arrowheads**). This is consistent with results showing that MyoF binds half as many calmodulin light chains as a construct containing a myosin Va lever arm (**Fig 1I-K**). The C-terminal region of MyoFΔtail forms a globular structure approximately 10-12 nm is diameter (**Fig. 4A, green arrowheads, red**) rather than an extended coiled-coil rod like MyoF/Va (**Fig. 4B, green arrowheads, red**).

**Figure 4:**
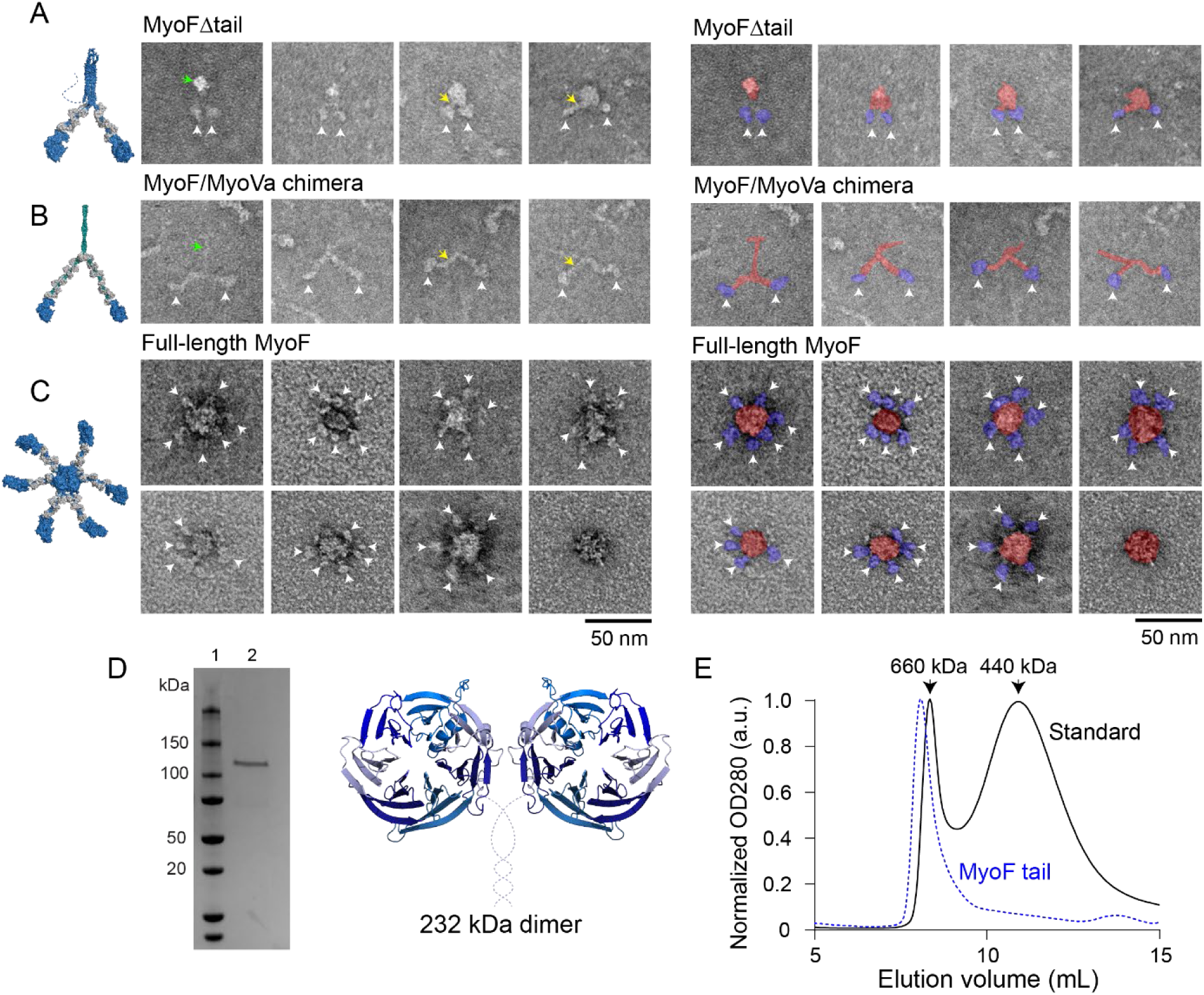
Electron micrographs and analytical SEC of MyoF constructs. Montage of negatively stained images of (**A**) MyoFΔtail co-purified with *Tg*Cam showing “v-shaped” particles typical of myosin dimers, (**B**) MyoF/Va chimeric construct, and (**C**) full-length MyoF, which appears as a central density associated with up to 6 motor domains. White arrowheads indicate MyoF motor domains, which are paired in MyoFΔtail and MyoF/Va chimeric constructs. Yellow arrowheads indicate the lever arm, which is approximately half the length for MyoF compared to the lever arm of mammalian myosin Va sequences. The green arrowhead points to the C-terminal dimerization domain which is compact for MyoF and an extended rod for MyoF/Va. Images are pseudo-colored (right) to indicate the motor domain (blue) and C-terminal regions (red). (**D**) Coomassie-stained SDS-PAGE gel (left) of the MyoF cc + WD40 construct (a.a. 987 – 1953) purified from *Sf9* cells, fused to a C-terminal biotin-FLAG tag. Cartoon depiction (right) of the MyoF cc + WD40 construct represented as a 232 kDa dimer. Dotted lined indicates sequence of unknown structure. (**E**) Analytical SEC showing the elution of 440 kDa (Ferritin, 10.98 mL peak) and 660 kDa (Thyroglobulin, 8.37 mL peak) standards (black) compared to the elution of MyoF cc + WD40 construct (blue, 8.18 mL peak) indicating that the WD40 tail domain oligomerizes into a complex that is at least hexameric.

To determine how the WD40 tail domain affects the organization of MyoF, we purified the full-length motor containing a C-terminal biotin-FLAG tag co-expressed with *Tg*Cam (**Fig. S2E**) and visualized negatively stained particles using EM. The full-length protein contains a central circular density approximately 20-25 nm in diameter, in most cases with associated multiple densities approximating the size of myosin motor domains. Particles contained up to six head domains, which may indicate a hexameric organization (**Fig. 4C)**.

Negatively stained images indicate that the WD40 tail domain of MyoF is capable of forming a higher order oligomer. To test this, we purified a MyoF tail construct containing the coiled-coil region, linker, and WD40 domain (a.a. 987 – 1,953) (MyoF cc + WD40) (**Fig. 4D**). When applied to a calibrated SEC column, the protein eluted before the 660 kDa standard (**Fig. 4E**). Because the molecular weight of a MyoF cc + WD40 dimer is 232 kDa, this indicates that the oligomerization state is at least hexameric (696 kDa). Because a protein of this molecular weight would elute in the void volume, it is not possible to distinguish a hexamer from a higher order oligomer. Taken together, these data indicate that the WD40 tail changes the stoichiometry from a two-headed dimer into a larger complex containing at least six heads.

### Full-length MyoF is processive on skeletal actin bundles and TgAct1 filaments

Because full-length MyoF oligomerizes into a multi-headed complex, it may move processively on actin as a single molecule. To test this, we imaged single molecules of full-length MyoF moving on skeletal actin filaments. Full-length MyoF containing a C-terminal biotin tag co-purified with *Tg*Cam was clarified and labeled with Alexa Fluor 647 streptavidin at a 1:5 molar ratio to ensure the fluorescent dye binds to a single motor. We find that single motors did not move processively on skeletal actin filaments (**Movie S9**).

The motility of small ensembles of MyoFΔtail bound to a Qdot was enhanced on actin bundles, so we next wondered if full-length MyoF was processive on bundles actin. To test this, single molecules of full-length MyoF co-purified with *Tg*Cam were imaged moving on actin-fascin bundles which were labeled and stabilized with Alexa Fluor 488-phalloidin. We observed many static associations of the full-length motor with these bundles (**Fig. 5A, left, vertical lines**). A few motors moved however, motility was poor, with frequent pausing and slow run speeds (**Fig. 5A-B, left**), giving rise to short run lengths (**Fig. 5C**).

**Figure 5:**
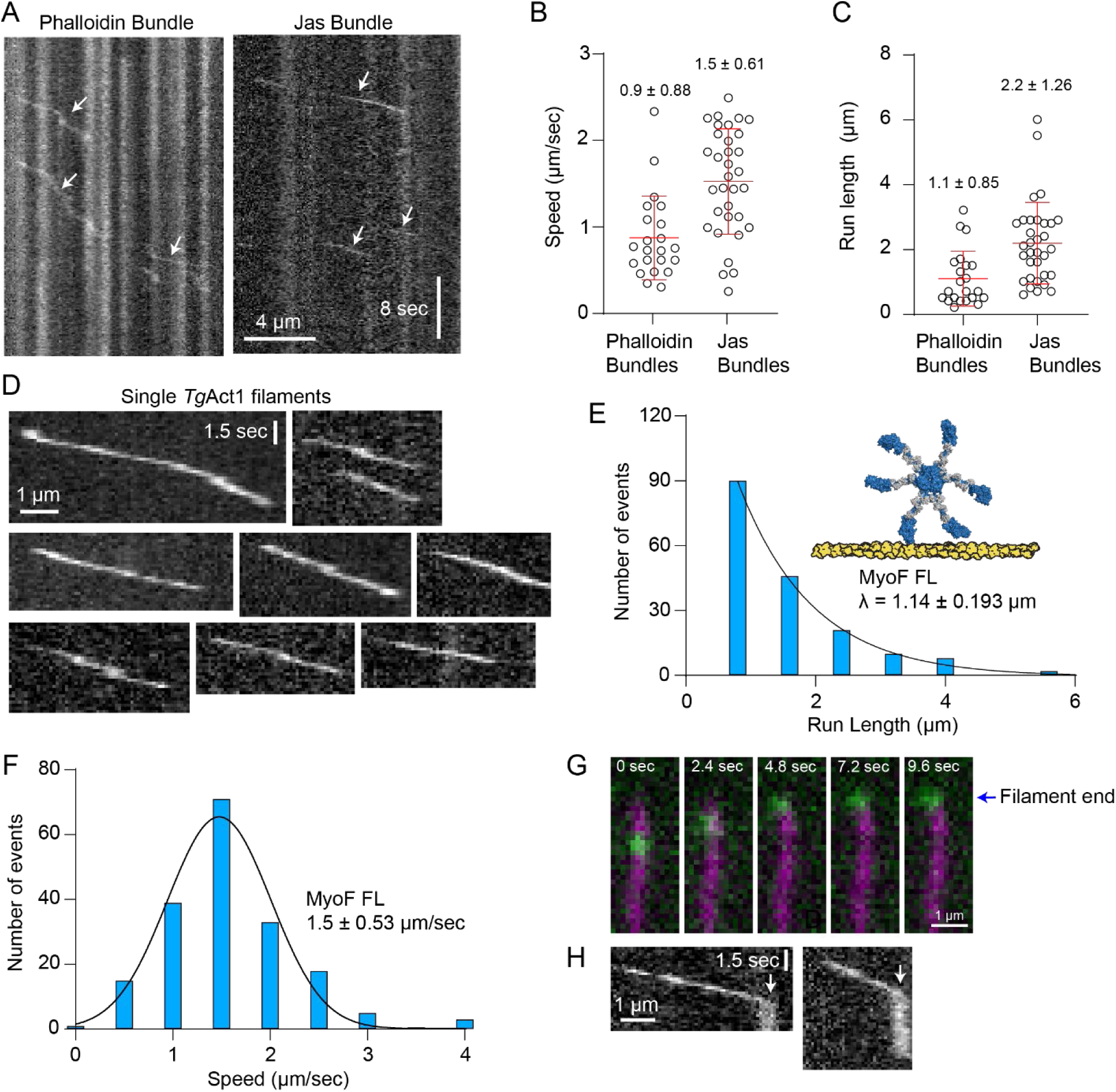
Motile properties of full-length MyoF. (**A**) Kymographs (distance versus time) of single molecules of full-length MyoF bound to *Tg*Cam labeled with Alexa Fluor 647 streptavidin moving on actin-fascin bundles made from skeletal actin stabilized with (left) phalloidin or (right) jasplakinolide. (**B**) Speed and (**C**) run length distributions for single molecules of full-length MyoF moving on actin-fascin bundles from skeletal actin stabilized with either phalloidin or jasplakinolide. Mean ± SD is shown (jasplakinolide bundles, n = 33; phalloidin bundles, n = 22). Average run speeds are significantly different (P value < 0.0001, using a two-tailed t-test). Average run lengths are significantly different (P value = 0.0008). (**D**) Kymogrpahs of single molecules of full-length MyoF labeled with Alexa Fluor 647 streptavidin moving on jasplakinolide stabilized *Tg*Act1 filaments. (**E**) Run length distribution (characteristic run length distribution (λ) ± error of the fit shown) and (**F**) speed distribution (average ± SD shown) of full-length MyoF bound to *Tg*Cam moving on *Tg*Act1 filaments (n = 187). (**G**) Montage showing full-length MyoF bound to Alexa Fluor 647 streptavidin (green) moving to the end of a TgAct1 filament (magenta). The *Tg*Act1 filament was imaged using the chromobody fused to EmeraldFP. (**H**) Kymographs of MyoF showing movement on a *Tg*Act1 filament (diagonal line), followed by retention at the end of filaments (vertical line, white arrow).

A structural analysis of rabbit skeletal muscle α-actin found that actin stabilizing agents can trap actin in different structural states. These studies show that the D-loop of actin bound to phalloidin adopts a closed conformation while filaments bound to jasplakinolide induces an open D-loop conformation (39). Our recent study showed that unlike skeletal actin, the D-loop of unstabilized *Tg*Act1 filaments adopt an open D-loop conformation (40) and we hypothesized that full-length MyoF would move more efficiently on actin fascin bundles stabilized with jasplakinolide, which might better mimic *T. gondii* actin filaments. To test this, jasplakinolide stabilized skeletal actin was bundled with fascin and imaged in flow chambers with actin chromobody fused to EmeraldFP, which we previously used to image real time *Tg*Act1 assembly *in vitro* (40). Single molecules of labeled full-length MyoF moved efficiently on these bundled actin structures with minimal pausing, faster speeds and long run lengths (**Fig. 5A-C, right**).

Next, we determined if full-length MyoF was processive on single *Tg*Act1 filaments. Alexa Fluor647 streptavidin-labeled full-length MyoF was added to flow chambers adsorbed with jasplakinolide stabilized *Tg*Act1 filaments which were visualized using the actin chromobody-EmeraldFP. We find that unlike skeletal actin, single molecules of full-length MyoF moved processively on *Tg*Act1 filaments (**Fig. 5D, Movie S10**) with a characteristic run length (λ) of 1.14 µm and average speed of 1.5 µm/sec (**Fig. 5E-F**). Interestingly, we often see MyoF retained at the end of the filament before dissociating (**Fig. 5G**). This feature is not typical of cargo transporting myosin’s and may provide additional insight into its cellular function. To determine how *Tg*Mlc1 influences the processivity of MyoF, we purified the full-length motor containing a C-terminal biotin tag co-expressed with *Tg*Cam and *Tg*Mlc1 (**Fig. S2F**). We find that when bound to both light chains, full-length MyoF was retained to the ends of many filaments, but it did not support processive movement on *Tg*Act1 (**Movie S11**).

To gain deeper mechanistic insight into why full-length MyoF is processive on *Tg*Act1 filaments and not skeletal actin filaments, we compared the movement of a MyoF/Va chimeric construct (which contains the MyoF motor domain and myosin Va backbone) (Fig. 1I) on skeletal and *Tg*Act1 actin isoforms. The use of chimeric myosin constructs was previously used by Krementsova *et. al.* to define the structural and functional features that contribute to processive movement of class V myosins (41). We find that multiple MyoF/Va motors bound to a Qdot did not move on skeletal actin (**Movie S12**) but runs were observed on jasplakinolide-stabilized *Tg*Act1 actin filaments (**Movie S13**). The characteristic run length (λ = 0.45 µm) of MyoF/Va (**Fig. S4A**) was much shorter compared to full-length MyoF (**Fig. 5E**) and the average speed of movement was similar to MyoFΔtail bound to *Tg*Cam (2.6 µm/sec) (**Fig. S4B**). We also find that unlike the full-length motor, the MyoF/Va dimer is non-processive as a single motor on *Tg*Act1 filaments (**Movie S14**).

### MyoF binds microtubules via its tail domain and localizes to intraconoid microtubules in vivo

Next, we sought to gain further insight into MyoF’s mechanism of action in the cell. We previously showed that MyoF localizes to both the cytosol and the parasite periphery, with the cytosolic pool controlling the dynamics of cytoplasmic actin and the peripheral pool driving a peripheral actin flow (15, 16). First, we sought to determine how MyoF controls cytoplasmic actin organization. One hypothesis, motivated by the structural similarity between the MyoF WD40 domain and other actin binding proteins (coronin and Aip1), is that the MyoF tail may bind actin filaments and control actin organization by crosslinking actin filaments via its WD40 and motor domains. To investigate this, we tested whether the MyoF tail binds actin filaments. Recombinantly purified MyoF cc + WD40 was labeled with Alexa Fluor 488 streptavidin and was added to a flow cell bound with rhodamine-phalloidin stabilized skeletal actin filaments. Epifluorescence microscopy showed no association of the tail with single actin filaments, even under low salt conditions (**Fig. S5**). Thus, in light of our evidence showing that the MyoF dimer further oligomerizes by the WD40 tail domain, this result supports an alternative model where MyoF drives cytosolic actin dynamics by interacting with multiple filaments simultaneously through its N-terminal head domains rather than the tail to translocate filaments relative to one another.

Next, we sought to understand how MyoF is anchored to the parasite periphery. We find that tubulin consistently copurifies with full-length MyoF from both *Sf*9 and *T. gondii* lysates (**Fig. S2E-F, Fig. Table S1**). *T. gondii* has several tubulin-based structures including 22 subpellicular microtubules that run approximately two-thirds the length of the parasite, the conoid made up of 14 tubulin fibers, and two intraconoid microtubules (**Fig. 6A**) (42, 43). To test if MyoF associates with any tubulin-based structures, we used Ultrastructure Expansion Microscopy (U-ExM) to determine the precise peripheral localization of MyoF-GFP. Visualization of MyoF-GFP with acetylated tubulin, a marker for subpellicular microtubules and the conoid show that MyoF localizes to the periphery as discrete spots which do not co-localize with the subpellicular microtubules (**Fig. 6B, inset b**), which suggests association with another pellicular protein within the inner membrane complex (IMC) (44). However, closer inspection reveals that MyoF localizes as a fiber in the center of the conoid (**Fig. 6B, inset a**), consistent with localization to the intraconoid microtubules.

**Figure 6:**
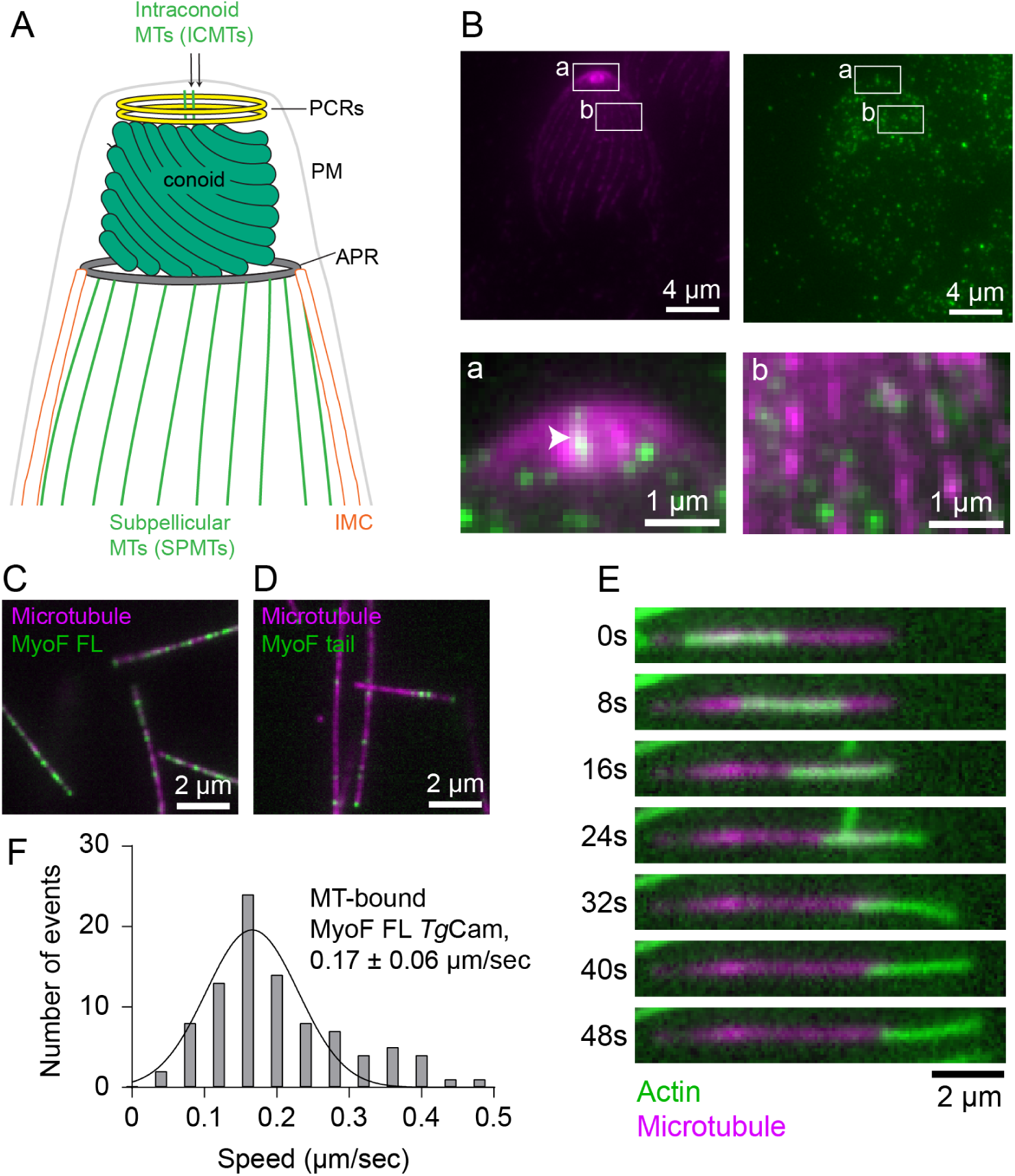
Cellular localization of MyoF by super resolution microscopy and translocation of actin on MyoF decorated microtubules. (B) Cartoon showing the cytoskeletal organization of the conoid, subpellicular microtubules, inner membrane complex (IMC), apical polar ring (APR), plasma membrane (PM) preconoidal rings (PCRs, yellow), and intraconoid microtubules (ICMTs, green) in *T. gondii*. (**B**) Ultrastructure Expansion Microscopy (U-ExM) showing the localization of acetylated tubulin (magenta) and MyoF-GFP (green). Merged insets show the filamentous localization of MyoF in the center of the conoid (inset a, arrowhead) and to the IMC (inset b). (**C-D**) Epifluorescence microscopy image showing that full-length MyoF-Alexa647 (B, green) and MyoF cc + WD40-Alexa647 (C, green) bind statically to microtubules (magenta). (**E**) Montage showing the movement of a skeletal actin filament (green) along a MyoF-bound microtubule (magenta). (**F**) Speed distribution of actin motility on MyoF bound microtubules (average speed ± SD, 0.17± 0.06. n = 94).

### MyoF translocates actin along microtubules

To determine if MyoF binds microtubules directly, we labeled the full-length motor with Alexa Fluor 647 streptavidin and added it to flow chambers containing fluorescently labeled taxol stabilized microtubules. We find that full-length MyoF bound to microtubules at nM concentrations (**Fig. 6C**). We also find that the MyoF cc + WD40 construct labeled with Alexa Fluor 647 streptavidin also binds microtubules, indicating that the interaction is medicated though its tail domain (**Fig. 6D**). Time lapse movies indicate that the binding for both constructs is static, suggesting that the binding is likely not a consequence of an electrostatic charge-charge interaction between the motor and microtubule E-hooks (**Movie S15-16**).

Because the tail binds to the microtubule lattice, the motor domains could be oriented to interact and move actin along the microtubule. To test this hypothesis, we decorated microtubules with full-length MyoF in flow chambers and added fluorescently labeled skeletal actin filaments. We find that the actin filaments associated with, and are translocated along, microtubules (**Fig. 6E**) (**Movie S17**) with an average speed of 0.17 µm/sec (**Fig. 6F**). Similar to the end retention observed in single molecule assays (**Fig. 5G-H**), filaments were often found to be retained at their plus (barbed)-end before dissociating (**Fig. 6E**).

## Discussion

Myosin F is a class 27 myosin motor that is unique to Apicomplexan parasites and protozoa in the alveolate group of eukaryotes (10). Loss of MyoF in *T. gondii*, results in defects in the organization and dynamics of the actin cytoskeleton which in turn disrupts vesicle transport, organelle inheritance during cell division and organization of the endomembrane pathway (4, 11, 12). To understand how MyoF functions in this process, we studied the properties of this motor which are completely undefined.

### MyoF forms an oligomer using its WD40 tail domain

The defining feature of MyoF is the presence of a WD40 tail domain (10). Approximately 1% of proteins in the human genome contain WD40 domains and are thus found in proteins with diverse cellular functions and often act as scaffolds for the formation of large protein complexes (45, 46). The WD40 domains in the actin binding proteins coronin and Aip1 bind to actin directly so we hypothesized that MyoF may regulate actin organization by crosslinking actin filaments via its motor domain and WD40 tail domain as several other molecular motors regulate the organization of their respective cytoskeletal tracks in this manner. A myosin motor complex containing Myo51 and associated proteins Rng8/9 from fission yeast, organizes the actin cytokinetic ring in fission by crosslinking and sliding actin filaments using actin binding sites in the motor domain and tail (47, 48). The mitotic kinesin Eg5 is a tetramer and slides spindle microtubules using pairs of motor domains at opposing ends of the molecule. Our data shows that the WD40 domain of MyoF does not bind single filaments, rather the full-length MyoF forms an oligomer consisting of a central hub formed from its WD40 domains. Emanating from this, we visualized up to six densities that were the size of myosin motor domain. Since the coiled-coil domain of MyoF can induce dimerization of the MyoFΔtail construct, this suggests that full-length MyoF forms a hexamer, or trimer of dimers. While speculative, particles with fewer than six heads visible could result from autoinhibitory head to tail interactions that promote a compact organization and prevent visualization of all motor domains. Formation of a multi-motor domain complex would allow the motor to associate with adjacent actin filaments and slide them relative to one another thereby controlling actin organization and dynamics in the cell (**Fig. 7, inset 1**).

**Figure 7:**
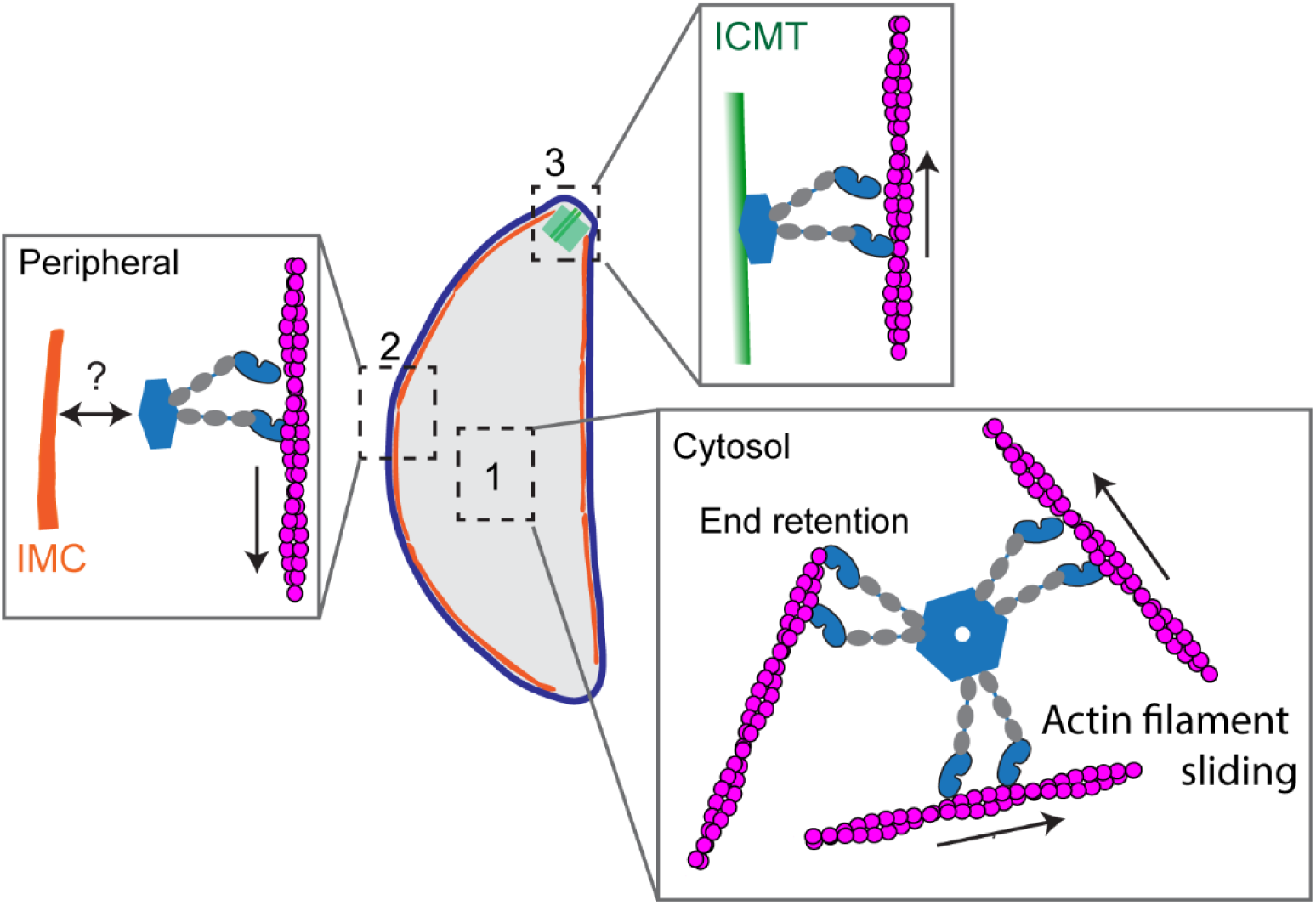
Model of MyoF’s mechanism of action. MyoF has three subcellular localizations. *Inset 1:* Hexameric MyoF organizes cytoplasmic actin filaments. Multiple pairs of motor domains interact with and translocate actin filaments thereby driving their dynamics in the cell. *Inset 2*: MyoF associates with the inner membrane complex through an unknown binding adapter. *Insert 3*: MyoF localizes to the intraconoid microtubules. The function of MyoF at the conoid requires further investigation. In inserts 2 and 3, the oligomeric state of MyoF at these localizations is unknown and is drawn as a dimer for simplicity.

### MyoF WD40 domain binds microtubules

In cells, MyoF associates with the parasite periphery, where we hypothesize it drives peripheral actin flow and movement of dense granules adjacent to the pellicle (4, 15). No colocalization between MyoF and the subpellicular microtubules was observed indicating that peripheral anchoring is likely driven by an, as yet unidentified, IMC component (**Fig. 7, inset 2**). MyoF did have an apical localization that was consistent with the intraconoid microtubules (49) (**Fig. 6B**). These microtubules are ∼350nm in length, run through the center of the hollow conoid and play a role in positioning the rhoptries inside the conoid (50). In addition, the ICMT’s associate with several microtubule-associated vesicles (MVs) of unknown function. Given the role of MyoF in the transport of other vesicle types (including dense granules and, Rab6- and ROP1-positive vesicles) it will be of interest in future studies to determine if MyoF has a role in positioning or transport of the rhoptries or MVs within the conoid (**Fig. 7, inset 3**). Association between MyoF and the ICMT’s is likely direct as full-length and tail constructs exhibited robust binding to taxol-stabilized brain microtubules at nM concentrations. Full-length MyoF could crosslink and slide actin filaments in relation to microtubules. Thus, in the cell, IMC and ICMT associated MyoF could drive actin filament dynamics in their respective cellular locations.

### MyoF motility is influenced by the composition of the actin track

Single molecules of MyoFΔtail (dimers) and full-length MyoF (oligomers) are non-processive on phalloidin-stabilized single filaments skeletal actin. While small ensembles of MyoFΔtail motors move inefficiently (i.e. - short run-lengths and a low run frequency). In contrast, full-length MyoF moved processively on jasplakinolide stabilized single *Toxoplasma* filaments and exhibited longer run-lengths and faster speeds on jasplakinolide stabilized skeletal bundles compared with phalloidin stabilized bundles (**Fig. 5A-C**) indicating that motor activity is influenced by the structural states of F-actin. While Toxoplasma and skeletal actin have similar overall helical parameters, we previously found an altered D-loop confirmation in *Toxoplasma* actin compared to skeletal actin (40). Jasplakinolide and phalloidin stabilized skeletal actin exhibit an “open” D-loop confirmation (39). Since the motile properties of other molecular motors are influenced by changes at the D-loop (51), it is tempting to speculate that it is this modification that confers track specificity of MyoF, however future structural studies will be required to define the basis of the motors altered activity. Taken together, these data indicate that the enhanced motility on *Tg*Act1, which shares only 83% identity to skeletal actin filaments, results from a preference of the MyoF motor domain with its native track. Because a two-headed MyoF construct is non-processive on *Tg*Act1 filaments, the duty cycle of MyoF is insufficient to support motility of a two-headed MyoF construct on *Tg*Act1 filaments. Thus, oligomerization by the MyoF cc + WD40 is necessary to form a ∼6-headed complex that supports single molecule motility of the full-length motor.

MyoF motility was enhanced on actin bundles compared with single filaments, with an ∼300-fold increase in run frequency (**Fig. 3C**). In addition, full-length accumulates at the tips of filopodia in *Sf*9 cells (**Fig. S3**). Several classes of myosin motors are optimized for movement on bundles including MyoVc (38) and Myo10 (29, 30, 52), although the actin bundle preference of these motors is conferred via different mechanisms. MyoVc forms a parallel dimer through is coiled-coil domain and additional flexibility in the lever arms accommodates bundle motility (38), while MyoX forms an anti-parallel coiled-coil so the orientation of the heads is optimized for movement on adjacent filaments on actin bundles in filopodia (31). In both these cases, motors encounter actin filament bundles in cells (53, 54). The physiological implications of MyoF’s enhanced motility on bundles requires further investigation because it is not known if *Toxoplasma* actin forms bundled networks in cells. The high turnover rate of *T. gondii* actin (40) precludes imaging in fixed cells (16), and makes ultrastructural imaging a challenge. In addition, *T. gondii* does not contain a fascin homolog. In fact, no actin bundling proteins have been identified in *T. gondii* to date so whether MyoF encounters parallel actin filament bundles in cells remains to be determined.

### Myosin light chain composition regulates MyoF velocity

MyoF contains 6 identifiable IQ motifs based on primary amino acid sequence. However, negative stain EM and a calmodulin binding assay showed that MyoF has a short lever arm that binds only three light chains. This is reminiscent of other myosins where there is a mismatch between the number of IQ motifs and number of associated light chains. *Leishmania* Myosin-XXI has 16 predicted light chain binding IQ motifs but only binds a single calmodulin (26), and the third IQ sequence in Myosin X does not bind a light chain, collectively indicating that presence of an identifiable IQ motif is not a reliable predictor of light chain binding (55).

We used immunoprecipitation to identify *Tg*Cam and *Tg*Mlc1 as the light chains associated with MyoF. *Tg*Cam binding is consistent with a previous report that identified MyoF as a close interacting partner of *Tg*Cam (56). *Tg*Mlc1 is a known binding partner of another parasite specific myosin (MyoA) involved in parasite motility and invasion (57). Data from the gliding filament assays (**Fig. 1**) and myosin motility assays (**Fig. 2**) showed that MyoF velocities is dependent on light chain composition. Because MyoFΔtail bound to *Tg*Cam supports *in vitro* gliding motility, we assume that the lever arm is fully occupied. While in the presence of *Tg*Mlc1 alone MyoF is non-motile indicating incomplete occupancy. When binding both *Tg*Cam and *Tg*Mlc1, MyoF velocity is ∼ 6 fold slower than when the motor is bound to *Tg*Cam alone indicating that *Tg*Mlc1 serves a regulatory function.

Despite the importance of the actin cytoskeleton to *T. gondii* growth and pathogenicity, the proteins and mechanisms which regulate actin organization are poorly understood. Characterizing MyoF’s unique domain organization and biophysical properties revealed a novel myosin-based mechanism of actin organization and dynamics. Given the absence of canonical proteins that control the formation of bundled and branched actin networks in *T. gondii* and related organisms, motor-driven actin organization may represent an evolutionary divergent mechanism of actin regulation.

## Methods

### Identification of MyoF-associated light chains

To identify MyoF associated light chains, ∼1x10^9^ extracellular MyoF-EmFP parasites (4) were lysed in 10 mL of GFP-trap buffer (25 mM Imidazole pH 7.4, 300 mM KCl, 1 mM EGTA, 0.5% TritonX-100, 2 mM DTT and 1:100 protease inhibitor cocktail (Millipore Sigma: Cat # P8340)) on ice for 10 minutes. Lysates were precleared at 16,000 g for 20 min at 4°C. 25 µL of GFP-Trap (ProteinTech Cat #GTMA) were resuspended and then washed x 2 in GFP-trap buffer. The cleared lysates were added to the magnetic beads and incubated with gentle shaking for 1 hour at 4°C. Using a magnetic rack, beads were washed ten times in GFP-trap buffer. After the final wash beads were wash x 2 in 1x PBS and resuspended in a final volume of 100µL. 10µL of beads were removed and added to an equal volume of 2x SB and 50 mM DTT, boiled for 5 minutes at 95°C. The remaining beads were stored in 1x PBS. LC MS/MS was performed at the University of Connecticut Proteomics and Metabolic Facility.

### Constructs

*Tg*MyoF (ToxoDB: TgME49_278870) coding sequence was cloned via PCR using RH cDNA. MyoF was then subcloned into a pFastBac expression vector so that the protein would be expressed in frame with a C-terminal FLAG tag used for affinity purification followed by a biotin tag for conjugation to fluorescent streptavidin reporters. (**Fig. S1B**). The 88-amino acid biotin tag is derived from the *Escherichia coli* biotin carboxyl carrier protein which becomes biotinylated at a single lysine residue (22, 23). For purification from bacteria, *Tg*Mlc1 containing an N-terminal 6xHIS tag was cloned into the pET3a bacterial expression vector and *Tg*Cam containing a C-terminal 6xHIS tag was cloned into the pET22b bacterial expression vector. For co-expression in *Sf*9 cells, untagged *Tg*Mlc1 and *Tg*Cam were cloned into pFastBac for baculovirus expression. The sequence of human fascin (FSCN1) was codon optimized for *E. coli* was fused at its N-terminus with a HAT-mNeonGreen sequence for HIS affinity purification and visualization using fluorescence microscopy. A TEV cleavage site was inserted between mNeon and FSC1 to generate untagged fascin. This sequence was inserted into pET16b for expression in BL21 (DE3) *E. coli*.

### Protein expression and purification

*T. gondii* light chains *Tg*Cam and *Tg*Mlc1 were fused to a 6xHIS tag for nickel affinity purification from BL21 (DE3) bacterial cells. Volumes of 500 mL Luria Broth (LB) were inoculated with a 25 mL starter culture of BL21 (DE3) transformed with *Tg*Cam or *Tg*Mlc1 light chain expression constructs and grown to an optical density of ∼0.8-1.2. Cells were then chilled, induced for protein expression with the addition of 0.4 mM isopropyl β-D-1-thiogalactopyranoside (IPTG), and incubated in a 16°C shaking incubator overnight. Cells were then harvested by centrifugation and frozen at -20°C. Cell pellets were thawed and resuspended in 50 mL ice cold lysis buffer (10 mM NaPO_4_, 300 mM NaCl, 0.5% glycerol, 7% sucrose, 0.5 mM PMSF, 0.5 mM DTT, 0.5% NP-40, protease inhibitors (Pierce Cat # A32953), pH 7.4) and lysed by incubating with 3 mg/mL lysozyme (Sigma Cat # L6876) on a 4°C rocking incubator for 30 minutes. Lysed cells were then sonicated and centrifuged for 35 min at 250,000 *g*. Supernatant was added to 1 mL prepared nickel select affinity resin (Cat # P6611) equilibrated with 0.5 M Imidazole followed by wash buffer and batch incubated at 4°C on a rocking platform for 45 minutes. The lysate was then applied to a chromatography column and washed with 60 column volumes of HIS wash buffer (10 mM NaPO_4_, 300 mM NaCl, 0.5% glycerol, 10 mM Imidazole, pH 7.4), followed by 20 column volumes HIS wash buffer containing 10 mM Imidazole. Bound protein was then eluted with elution buffer (10 mM NaPO_4_, 300 mM NaCl, 0.5% glycerol, 200 mM Imidazole, pH to 7.4, 0.5 mM DTT) in 1 mL fractions. Fractions that were positive for protein by Bradford were pooled, concentrated to ∼2 mL and dialyzed against storage buffer (25 mM Imidazole, pH 7.4, 300 mM NaCl, 50% glycerol, 1 mM DTT, 1 µg/mL leupeptin) overnight at 4°C.

Human fascin was purified similarly to light chains except, following elution from nickel select affinity resin, ∼10 mg of protein was dialyzed into TEV cleavage buffer (10 mM phosphate, pH 7.4, 50 mM NaCl, 1 mM DTT), clarified 400,000 g for 15 min at 4°C, and cleaved with TEV protease for 1 hour at 27°C. NaCl was then added to the mixture at a final concentration of 250 mM and 0.25 mL of nickel select affinity resin equilibrated in 0.5 M Imidazole, followed by TEV binding buffer was added and incubated on a nutating rocker at 4°C for 1 hour. The mixture was then centrifuged and the supernatant containing untagged fascin was dialyzed in storage buffer (25 mM Imidazole, pH 7.4, 300 mM NaCl, 50% glycerol, 1 mM DTT, 1 µg/mL leupeptin) overnight at 4°C.

Myosin constructs were co-expressed with the indicated light chain/s in *Sf*9 cells using the baculovirus system. Infected *Sf*9 cells were incubated for 72 hours at 27°C, harvested by centrifugation and resuspended in 60 mL FLAG lysis buffer (10 mM Imidazole, pH 7.4, 0.3 M NaCl, 5 mM MgCl_2_, 7% sucrose, 1 mM EGTA, 1 mM DTT, 2 mM MgATP, 25 µg/ml indicated light chain, and protease inhibitors (1x Sigma (Cat # P8340), 0.5 mM PMFS, 5 µg/ml leupeptin (Sigma Cat # L2884, and 1 mg/ml benzamidine). Cells were lysed by sonication and then clarified by centrifugation for 35 min at 250,000 *g*. The supernatant was added to 2 mL FLAG affinity resin (ThermoScientific Cat # PIA36803) equilibrated in FLAG wash buffer (10 mM imidazole, pH 7.4, 0.3 M NaCl, and 1 mM EGTA) and incubated in batch for 45 minutes on a rocking shaker at 4°C. The FLAG resin was then collected by gentle centrifugation (5 min at 800 rpm) and applied to a column for washing with ∼60 mL FLAG wash buffer at 4°C. Bound protein was then eluted with 0.1 mg/mL FLAG peptide in wash buffer. Fractions positive for protein by Bradford were pooled and concentrated to 0.5 mL using a 10 kDa MWCO Amicon Ultra Centrifugal Filter (Millipore, Cat # 2022-08-01) and applied to a Superdex 200 Increase 10/300 GL (GE Healthcare, 28990944) gel filtration column equilibrated in FLAG wash buffer. Fractions corresponding to full-length MyoF were pooled, concentrated to ∼1.5 mL, and dialyzed against storage buffer (25 mM Imidazole, pH 7.4, 300 mM NaCl, 50% glycerol, 1 mM DTT, 1 µg/mL leupeptin) overnight at 4°C.

Skeletal actin, *T. gondii* actin, actin chromobody (actin CB-EmFP), and NEM myosin were prepared as described in our previous publication (40). Kinesin406 G235A (“rigor” kinesin) was prepared as described previously (58).

### In vitro gliding filament assay

*In vitro* motility flow chambers were coated with a nitrocellulose film and adsorbed with biotinylated bovine serum albumin (Bio-BSA) by incubating with 0.5 mg/mL Bio-BSA for 1 minute in buffer B150 (150 mM KCl, 25 mM Imidazole, pH 7.4, 1 mM EGTA, 4 mM MgCl_2_, and 10 mM DTT). Flow cells were then washed and blocked with 0.5 mg/ml BSA in buffer B150 for 1 minute, washed and functionalized with neutravidin by adding 50 μg/mL neutravidin (Thermo Fisher Scientific) in buffer B150 for 1 minute followed by three washes with buffer B150 to remove unbound neutravidin. MyoF constructs were prepared by performing an initial actin spin-down to remove motors that are unable to dissociate from actin in the presence of MgATP. To do this, MyoF constructs were diluted to 0.7 μM in buffer B150. To this 1.4 μM skeletal F-actin and 10 mM MgATP was added, mixed and centrifuged 20 min at 350,000 g. The soluble fraction was then applied to the neutravidin functionalized flow chamber for 2 minutes and washed with buffer B150 to remove unbound motor and MgATP. The flow chamber was then infused with 50 nM rhodamine-phalloidin-labeled actin for 1.5 minutes and washed with buffer B (50 mM KCl, 25 mM Imidazole, pH 7.4, 1 mM EGTA, 4 mM MgCl_2_, and 10 mM DTT). Gliding motility was initiated with the addition of In vitro motility Go buffer (50 mM KCl, 25 mM Imidazole, pH 7.4, 1 mM EGTA, 4 mM MgCl_2_, 0.25% methylcellulose, 10 mM DTT, 0.1 mg/mL of the indicated light chain, 2 mM MgATP, an ATP regeneration system (1 mM phosphocreatine, 0.1 mg/mL creatine phosphokinase) and oxygen scavenging system (50 μg/mL catalase, 3 mg/mL glucose, and 125 μg/mL glucose oxidase). Gliding filaments were imaged at 37 °C in epifluorescence using a DeltaVision Elite microscope (Cytiva) built on an Olympus base with a 100 × 1.39 NA objective and definite focus system. Images were acquired every 1 second for 2 minutes with a scientific CMOS camera and DV Insight solid-state illumination module.

### Single and multiple motor motility assays

For imaging the motility of single molecules and small ensembles of MyoFΔtail constructs, the motor was diluted in buffer M (10 mM Imidazole pH 7.4, 4 mM MgCl_2_, 1 mM EGTA, 300 mM KCl) and clarified 400,000 g for 15 min at 4°C, and its concentration was determined by a Bradford assay. For single molecule experiments, 80 nM MyoFΔtail was mixed with 400 nM Qdot 655 streptavidin conjugate (1:5 ratio) in buffer M containing 4 mg/mL BSA and 0.5% Pluronic F-127.

At this ratio, the majority of Qdots are bound to a single motor. For experiments visualizing the motility of small ensembles of MyoFΔtail constructs, 1 μM MyoFΔtail was mixed with 0.1 μM Qdot 655 streptavidin conjugate (10:1 ratio) in the same buffer, which promotes the binding of multiple motors per Qdot. For the attachment of stabilized F-actin filaments, NEM-treated muscle myosin was absorbed to the surface of a flow chamber, washed with buffer B (50 mM KCl, 25 mM Imidazole, pH 7.4, 1 mM EGTA, 4 mM MgCl2, and 10 mM DTT) and blocked with blocking buffer (50 mM KCl, 25 mM Imidazole, pH 7.4, 1 mM EGTA, 4 mM MgCl_2_, 4 mg/mL BSA, 0.5% Pluronic F-127, and 10 mM DTT). The flow chamber was then rinsed with buffer B and infused with 50 nM rhodamine phalloidin or Alexa Fluor 488 phalloidin stabilized skeletal F-actin filaments. After 2 minutes, the flow chamber was rinsed with buffer B to remove unbound filaments, and myosin mixtures were diluted 1:50 to 1:200 into Go buffer (50 mM KCl, 25 mM Imidazole, pH 7.4, 1 mM EGTA, 4 mM MgCl_2_, 10 mM DTT, 4 mg/mL BSA, 0.5% Pluronic F-127, 2 mg/mL kappa casein, 0.1 mg/mL of the indicated light chain, 2 mM MgATP, and oxygen scavenging system (50 μg/mL catalase, 3 mg/mL glucose, and 125 μg/mL glucose oxidase)), added to the flow chamber, and imaged at 37 °C with an imaging rate of 0.05 to 0.2 seconds where indicated. Images were acquired using epifluorescence microscopy on a DeltaVision Elite microscope or using Total Internal Reflection Fluorescence (TIRF) microscopy on a Nikon AXR built around the inverted Ti2E microscope and captured on Photometrics Prime 95B sCMOS camera where indicated.

Single molecule experiments with full-length MyoF were done similarly except MyoF was prepared by diluting into B500 (500 mM KCl, 25 mM Imidazole, pH 7.4, 1 mM EGTA, 4 mM MgCl_2_, and 10 mM DTT) and centrifuged 400,000 *g* for 15 min at 4°C. The motor was then diluted to 300 nM and added to Alexa Fluor 647 streptavidin conjugate that was diluted into B500 and clarified 400,000 *g* for 15 min at 4°C at a molar ratio of 1:5. This mixing ratio ensures that the majority of streptavidin fluorophores are bound to a single motor. The motility of single full-length motors was visualized on Alexa Fluor 488 phalloidin stabilized skeletal F-actin bound to partially PEGylated flow cells (59). For single and ensemble experiments using *T. gondii* actin filaments, *T. gondii* actin filaments were stabilized with jasplakinolide at a molar ratio of 1:1.2 and attached to the surfaces of flow cells similarly to stabilized skeletal actin and imaged in Go buffer supplemented with 1 μM jasplakinolide and 50 nM actin chromobody-EmeraldFP for visualization.

### Dynamic actin-microtubule crosslinking assay

Labeled microtubules were prepared by polymerizing a mixture of cycled bovine tubulin and Cy5 tubulin (PurSolutions) followed by stabilization with paclitaxel (Cytoskeleton, Denver, CO) as described previously (58). To attach microtubules, kinesin406 G235A (“rigor” kinesin) was diluted to 0.1 mg/mL in buffer B and added to a partially PEGylated flow chamber for 2 minutes. The flow chamber was then washed with buffer B, blocked with blocking buffer for 2 minutes, and rinsed with buffer B. The flow chamber was then infused with 0.9 μM taxol stabilized Cy5-labeled microtubules for 2 minutes and then rinsed with buffer B containing 10 μM paclitaxel to remove unbound microtubules. Unlabeled clarified full-length MyoF was diluted to 10 nM in actin/MT Go buffer (80 mM KCl, 25 mM Imidazole, pH 7.4, 1 mM EGTA, 4 mM MgCl_2_, 10 mM DTT, 4 mg/mL BSA, 0.5% Pluronic F-127, 2 mg/mL kappa casein, 0.1 mg/mL *Tg*Cam, 0.25% methylcellulose, 10 μM paclitaxel, 2 mM MgATP, and oxygen scavenging system (50 μg/mL catalase, 3 mg/mL glucose, and 125 μg/mL glucose oxidase)) and added to the flow chamber for 5 minutes to allow the MyoF cc + WD40 to bind microtubules. The flow cell was then rinsed with buffer B containing 10 μM paclitaxel to remove unbound MyoF and then actin/MT Go buffer containing 0.1 μM Alexa Fluor 488 phalloidin stabilized skeletal F-actin was added to the flow chamber which was subsequently sealed using nail polish. Movement actin along microtubules was observed at 37 °C using epifluorescence microscopy and images were captured every 10 seconds for 20 minutes.

### MyoF cc + WD40-microtubule binding assay

The MyoF cc + WD40 (a.a. 987 – 1953) containing a C-terminal biotin-FLAG tag was diluted in buffer M, clarified 400,000 g for 15 min at 4°C, and its concentration was determined by a Bradford assay. The protein was diluted to 0.1 μM and mixed with 0.5 μM pre-clarified Streptavidin Alexa Fluor 488 Conjugate (ThermoFisher) to ensure binding of a single molecule per fluorophore. Partially PEGylated flow chambers were functionalized, passivated, and bound to fluorescently labeled microtubules as described above. The labeled MyoF cc + WD40 was diluted 1:25 in modified Go buffer (50 mM KCl, 25 mM Imidazole, pH 7.4, 1 mM EGTA, 4 mM MgCl_2_, 10 mM DTT, 4 mg/mL BSA, 0.5% Pluronic F-127, 2 mg/mL kappa casein, 10 μM paclitaxel, 1 mM MgATP, and oxygen scavenging system (50 μg/mL catalase, 3 mg/mL glucose, and 125 μg/mL glucose oxidase)), added to the flow chamber for 6 minutes, washed with modified Go buffer, and the microtubules and labeled MyoF were imaged using epifluorescence microscopy on a DeltaVision Elite microscope.

### MyoF cc + WD40 and full-length MyoF-actin binding assay

Rhodamine-phalloidin stabilized skeletal actin adhered flow chambers were prepared as described above. MyoF cc + WD40 in buffer M was and clarified 400,000 g for 15 min at 4°C, and its concentration was determined by a Bradford assay. The protein was then diluted to 1 μM and mixed with 0.1 Qdot 655 streptavidin conjugate (5:1 ratio) to promote the binding of multiple molecules per Qdot. The mixture was then diluted 1:50 in a modified low salt imaging buffer (10 mM KOAc, 25 mM Imidazole, pH 7.4, 1 mM EGTA, 4 mM MgCl_2_, 10 mM DTT, 0.5% Pluronic F-127, 2 mg/mL kappa casein, 2 mM MgATP, and oxygen scavenging system (50 μg/mL catalase, 3 mg/mL glucose, and 125 μg/mL glucose oxidase)), added to the flow chamber, and imaged using epifluorescence microscopy on a DeltaVision Elite microscope.

### Ectopic expression in Sf9 cells

20 million *Sf*9 cells grown to mid-log were infected with 1 mL of a baculovirus encoding full-length MyoF fused at the C-terminus the mClover3 variant of GFP and incubated 2.5 hours on a rocking platform at room temperature. Cells/virus mixture was then added to 5 mL of *Sf*9 media, and 2 mL was added to a MatTek dish and allowed to infect for 2 days at 25°C. Media was aspirated and 2 mL of cold 4% paraformaldehyde in 1x PBS was added to the well and fixed for 20 minutes. The well was then washed 3 times with 1x PBS, permeabilized with 0.25% TX-100, washed three times with 1xPBS. and 1x PBS containing 20 µM rhodamine phalloidin was added (ThermoFisher) and incubated on a nutator for 1 hour. The well was washed 3 times with 1x PBS and imaged in TRITC/FITC channels using epifluorescence microscopy on a DeltaVision Elite microscope.

### Expansion microscopy

Human foreskin fibroblasts (HFFs) were grown to confluency on 12 mm round coverslips in a 12 well plate and infected overnight with *T. gondii* parasites expressing endogenous levels of MyoF fused at the C-terminus to emeraldFP (MyoF-EmFP) (4). The following day, the cells were fixed in 4% paraformaldehyde at room temperature for 20 minutes, washed 3 times in 1xPBS, transferred to a 6 well plate containing FA/AA solution (1x PBS, 1.2% paraformaldehyde, 2% acrylamide), sealed with parafilm, and incubated at 37°C overnight. The next day, the FA/AA solution was removed, and cells were washed 1x with PBS. To induce polymerization, 90 μL monomer/gel solution (19% sodium acrylate, 10% acrylamide, 0.1% BIS, 1.1x PBS) was mixed with 5 μL of 10% TEMED and 5 μL of 10% ammonium persulfate (APS) and 35 μL of this solution was added to parafilm in a 6 well plate stored on ice. The coverslip was then immediately inverted onto the gel solution and allowed to polymerize for 5 minutes and then transferred to a 37°C incubator for 1 hour. Afterwards, 2 mL of denaturization buffer (50 mM Tris-HCl, pH 9, 200 mM sodium dodecyl sulfate (SDS), 200 mM NaCl) were added to the wells and placed on a rocker for 15 minutes at room temperature. Using forceps and a razer, the gel was detached from the coverslip and transferred to a 1.5 mL Eppendorf tube containing 1.5 mL of denaturization buffer and incubated for 90 minutes at 95°C. Gels were then transferred to a petri dish containing 25 mL of DI water and incubated for 30 minutes at room temperature. The expended gels were then washed twice with water, quartered with a razor and washed again with 1x PBS. One segment of the gel was transferred to a 12 well dish containing blocking buffer (2% BSA, 1x PBS, 0.5% Tween-20) and rocked on a shaker at room temperature for 30 minutes. The gel slice was treated with primary antibody (1:500 mouse anti-acetylated tubulin (Sigma T6793) and 1:500 rabbit anti-GFP-Alexa Fluor 488 (Invitrogen A21311)) diluted in blocking buffer and rocked overnight at room temperature. The next day, the gel was washed 3x in 2 mL wash buffer (1x PBS, 0.5% Tween-20) and bound to secondary antibody (1:500 goat anti-mouse Alexa Fluor 647 (ThermoFisher A-21235), 1:500 goat anti-rabbit Alexa Fluor 488 (ThermoFisher A-11008)) for 2.5 hours. The gel slice was then washed 3x with 2 mL wash buffer followed by 3x with distilled water incubating 30 minutes between each wash. The expanded gel was then transferred to a Poly-D-Lysine coated MatTek dish and imaged using epifluorescence microscopy on a DeltaVision Elite microscope. The expansion factor was determined by measuring the diameter of the gel before and after expansion.

## Acknowledgements

We thank Drs. Jeremy Balsbaugh and Jennifer Liddle at UConn Proteomics and Metabolomics for their assistance with the mass spectrometry analysis, Dr. Chris O’ Connell at the UConn Advanced Microscopy Facility for assistance with TIRF microscopy, Dr. Sabina Absalon (Indiana University), Dr. Christopher de Graffenried (Brown University), and Dr. Paul Campbell for assistance with the expansion microscopy experiments.

## Author Contributions Statement

TES and ATH contributed to the design and execution of experiments. Both authors co-wrote the paper.

## Tables

**Table 1.** Summary of MyoF interacting proteins from *T. gondii* cell lysates.

|  |  |  |
| --- | --- | --- |
| IP replicate 1 accession number | Annotation | # peptides |
| TGME49_278870 | myosin F | 118 |
| TGME49_239400 | inner membrane complex protein IMC28 | 13 |
| TGME49_227870 | Tim10/DDP family zinc finger superfamily protein | 3 |
| TGME49_257680 | myosin light chain MLC1 | 2 |
| TGME49_203260 | cell cycle checkpoint protein RAD17, putative | 1 |
| TGME49_203980 | hypothetical protein | 1 |
| TGME49_209910 | histone H2Bv | 1 |
| TGME49_221620 | beta-tubulin, putative | 1 |
| TGME49_223750 | hypothetical protein | 1 |
| TGME49_223940 | gliding-associated protein GAP45 | 1 |
| TGME49_228660 | Sec7 domain-containing protein | 1 |
| TGME49_228690 | phosphatidylinositol 3- and 4-kinase | 1 |
| TGME49_231630 | alveolin domain containing intermediate filament IMC4 | 1 |
| TGME49_232280 | hypothetical protein | 1 |
| TGME49_233460 | SAG-related sequence SRS29B | 1 |
| TGME49_254610 | Tim10/DDP family zinc finger superfamily protein | 1 |
| TGME49_259080 | hypothetical protein | 1 |
| TGME49_260790 | RAP domain-containing protein | 1 |
| TGME49_261250 | histone H2A1 | 1 |
| TGME49_264485 | AP2 domain transcription factor AP2IX-3 | 1 |
| TGME49_291050 | serine/threonine-protein kinase KIN | 1 |
| TGME49_293590 | 3-oxoacyl-acyl-carrier protein synthase I/II, putative | 1 |
| TGME49_308090 | roptry protein ROP5 | 1 |
| TGME49_309885 | hypothetical protein | 1 |
| TGME49_310010 | roptry neck protein RON1 | 1 |
| TGME49_311240 | DnaJ family chaperone J1 | 1 |
| TGME49_312520 | tRNA dimethylallyltransferase | 1 |
| TGME49_313140 | isocitrate dehydrogenase 2 | 1 |
| TGME49_316760 | hypothetical protein | 1 |

| IP replicate 2 accession number | Annotation | # peptides |
| --- | --- | --- |
| TGME49_278870 | myosin F | 302 |
| TGME49_249240 | calmodulin, putative | 45 |
| TGME49_290870 | patched family protein | 22 |
| TGME49_236650 | DEAD (Asp-Glu-Ala-Asp) box polypeptide 17 | 21 |
| TGME49_239400 | inner membrane complex protein IMC28 | 16 |
| TGME49_266960 | beta tubulin | 12 |
| TGME49_269190 | glyceraldehyde-3-phosphate dehydrogenase GAPDH2 | 11 |
| TGME49_273760 | heat shock protein HSP70 | 10 |
| TGME49_237180 | hypothetical protein | 10 |
| TGME49_233460 | SAG-related sequence SRS29B | 9 |
| TGME49_215775 | roptry protein ROP8 | 8 |
| TGME49_310010 | roptry neck protein RON1 | 8 |
| TGME49_232410 | thioredoxin-like protein 1 | 8 |
| TGME49_311720 | chaperonin protein BiP | 7 |
| TGME49_231630 | alveolin domain containing intermediate filament IMC4 | 7 |
| TGME49_235470 | myosin A | 7 |
| TGME49_316400 | alpha tubulin TUBA1 | 6 |
| TGME49_219320 | gliding-associated protein GAP50 | 6 |
| TGME49_231640 | alveolin domain containing intermediate filament IMC1 | 6 |
| TGME49_239260 | histone H4 | 6 |
| TGME49_257680 | myosin light chain MLC1 | 6 |
| TGME49_236540 | RNA recognition motif-containing protein | 6 |
| TGME49_286420 | elongation factor 1-alpha (EF-1-ALPHA), putative | 5 |
| TGME49_213392 | surface antigen repeat-containing protein | 5 |
| TGME49_319560 | microneme protein MIC3 | 4 |
| TGME49_258870 | cyst wall protein CST7 | 4 |
| TGME49_258580 | roptry protein ROP17 | 4 |
| TGME49_305160 | histone H2Ba | 4 |
| TGME49_223920 | roptry neck protein RON3 | 4 |
| TGME49_288650 | dense granule protein GRA12 | 4 |
| TGME49_313380 | IMC localizing protein ILP1 | 4 |
| TGME49_250340 | centrin 2 | 4 |
| TGME49_300200 | histone H2AZ | 4 |
| TGME49_289750 | ribosomal-ubiquitin protein RPL40 | 3 |
| TGME49_209910 | histone H2Bv | 3 |
| TGME49_261580 | histone H2AX | 3 |
| TGME49_311240 | DnaJ family chaperone J1 | 3 |
| TGME49_263090 | 14-3-3 protein | 3 |
| TGME49_249900 | adenine nucleotide translocator | 3 |
| TGME49_308090 | roptry protein ROP5 | 3 |
| TGME49_232940 | heat shock protein HSP20 | 3 |
| TGME49_203310 | dense granule protein GRA7 | 2 |
| TGME49_216000 | alveolin domain containing intermediate filament IMC3 | 3 |
| TGME49_279100 | mitochondrial association factor 1a | 3 |
| TGME49_295110 | roptry protein ROP7 | 3 |
| TGME49_218260 | histone H3.3 | 2 |
| TGME49_271050 | SAG-related sequence SRS34A | 2 |
| TGME49_295125 | roptry protein ROP4 | 2 |
| TGME49_209030 | actin ACT1 | 2 |
| TGME49_262620 | RNA recognition motif-containing protein | 2 |
| TGME49_211680 | protein disulfide isomerase | 2 |
| TGME49_248700 | alveolin domain containing intermediate filament IMC12 | 2 |
| TGME49_243950 | prohibitin, putative | 2 |
| TGME49_227280 | dense granule protein GRA3 | 2 |
| TGME49_261950 | ATP synthase beta subunit ATP-B | 2 |
| TGME49_205658 | F5/8 type C domain-containing protein | 2 |
| TGME49_201780 | microneme protein MIC2 | 2 |
| TGME49_231160 | hypothetical protein | 2 |
| TGME49_219270 | glideosome-associated protein with multiple-membrane spans GAPM2A | 2 |
| TGME49_207040 | hypothetical protein | 2 |
| TGME49_220270 | alveolin domain containing intermediate filament IMC6 | 2 |
| TGME49_235402 | CorA family Mg2+ transporter protein | 2 |
| TGME49_293590 | 3-oxoacyl-acyl-carrier protein synthase I/II, putative | 2 |
| TGME49_209485 | hypothetical protein, conserved | 1 |
| TGME49_207800 | hypothetical protein | 1 |
| TGME49_229480 | calcium binding protein precursor, putative | 1 |
| TGME49_215980 | hypothetical protein | 1 |
| TGME49_215980 | hypothetical protein | 1 |
| TGME49_217340 | hypothetical protein | 1 |
| TGME49_306030 | glutathione S-transferase 2 | 1 |
| TGME49_306920 | HOOK interacting protein HIP | 1 |
| TGME49_258410 | photosensitized INA-labeled protein PHIL1 | 1 |
| TGME49_208030 | microneme protein MIC4 | 1 |
| TGME49_235930 | domain K- type RNA binding proteins family protein | 1 |
| TGME49_257540 | hypothetical protein | 1 |
| TGME49_269980 | preprotein translocase Sec61, putative | 1 |
| TGME49_249480 | co-chaperone UNC | 1 |
| TGME49_288380 | heat shock protein HSP90 | 1 |
| TGME49_310460 | ras-related protein RAB6 | 1 |
| TGME49_219660 | hypothetical protein | 1 |
| TGME49_313140 | isocitrate dehydrogenase 2 | 1 |
| TGME49_305630 | F-box domain-containing protein | 1 |
| TGME49_306460 | bromodomain-containing protein BDP4 | 1 |
| TGME49_227080 | PCIF1 WW domain-containing protein | 1 |
| TGME49_229150 | transporter, major facilitator family protein | 1 |
| TGME49_204400 | ATPase synthase subunit alpha, putative | 1 |
| TGME49_272490 | protoporphyrinogen oxidase | 1 |
| TGME49_278130 | basal complex component BCC11 | 1 |
| TGME49_259630 | inner membrane complex protein IMC27 | 1 |
| TGME49_309885 | hypothetical protein | 1 |
| TGME49_224850 | polyadenylate-binding protein PABC | 1 |
| TGME49_270250 | dense granule protein GRA1 | 1 |
| TGME49_273445 | iron-sulfur cluster assembly protein SufD | 1 |
| TGME49_230160 | RNA-binding protein, putative | 1 |

| IP replicate 3 accession number | Annotation | # peptides |
| --- | --- | --- |
| TGME49_278870 | myosin F | 162 |
| TGME49_249240 | calmodulin, putative | 21 |
| TGME49_290870 | patched family protein | 11 |
| TGME49_236650 | DEAD (Asp-Glu-Ala-Asp) box polypeptide 17 | 9 |
| TGME49_239400 | inner membrane complex protein IMC28 | 5 |
| TGME49_237180 | hypothetical protein | 5 |
| TGME49_233460 | SAG-related sequence SRS29B | 4 |
| TGME49_215775 | roptry protein ROP8 | 4 |
| TGME49_310010 | roptry neck protein RON1 | 4 |
| TGME49_286420 | elongation factor 1-alpha (EF-1-ALPHA), putative | 4 |
| TGME49_226790 | ABC transporter, ATP-binding domain-containing protein | 2 |
| TGME49_305240 | XPA binding protein 2 family protein | 2 |
| TGME49_223985 | protein phosphatase PPM11B, putative | 2 |
| TGME49_292320 | queuine tRNA-ribosyltransferase, putative | 2 |
| TGME49_290620 | centrosomal protein CEP250L1 | 2 |
| TGME49_316400 | alpha tubulin TUBA1 | 1 |
| TGME49_219320 | gliding-associated protein GAP50 | 1 |
| TGME49_258580 | roptry protein ROP17 | 1 |
| TGME49_319560 | microneme protein MIC3 | 1 |
| TGME49_258870 | cyst wall protein CST7 | 1 |
| TGME49_289750 | ribosomal-ubiquitin protein RPL40 | 1 |
| TGME49_203310 | dense granule protein GRA7 | 1 |
| TGME49_271050 | SAG-related sequence SRS34A | 1 |
| TGME49_218260 | histone H3.3 | 1 |
| TGME49_207040 | hypothetical protein | 1 |
| TGME49_235402 | CorA family Mg<sup>2+</sup> transporter protein | 1 |
| TGME49_227620 | dense granule protein GRA2 | 1 |
| TGME49_234505 | phenylalanyl-tRNA synthetase alpha chain A, putative | 1 |
| TGME49_216170 | cysteine desulfurase NFS2 | 1 |
| TGME49_248160 | ATP-dependent DNA helicase 2 subunit KU70 | 1 |
| TGME49_212045 | hypothetical protein, conserved | 1 |
| TGME49_266880 | dihydrouridine synthase, putative | 1 |
| TGME49_255700 | hypothetical protein | 1 |
| TGME49_205470 | translation elongation factor 2 family protein, putative | 1 |
| TGME49_210700 | VPS13 domain-containing protein | 1 |
| TGME49_222930 | hypothetical protein | 1 |
| TGME49_223080 | hypothetical protein | 1 |
| TGME49_277950 | lipase | 1 |
| TGME49_237460 | DER1-like protein | 1 |
| TGME49_227010 | roptry kinase family protein ROP30 | 1 |
| TGME49_286450 | dense granule protein GRA5 | 1 |
| TGME49_244910 | MIZ/SP-RING zinc finger domain-containing protein | 1 |
| TGME49_240970 | hypothetical protein | 1 |

| Present in at least 2 of 3 datasets | Annotation | Total # peptides |
| --- | --- | --- |
| TGME49_278870 | myosin F | 582 |
| TGME49_249240 | calmodulin, putative | 66 |
| TGME49_239400 | inner membrane complex protein IMC28 | 34 |
| TGME49_290870 | patched family protein | 33 |
| TGME49_236650 | DEAD (Asp-Glu-Ala-Asp) box polypeptide 17 | 30 |
| TGME49_237180 | hypothetical protein | 15 |
| TGME49_233460 | SAG-related sequence SRS29B | 14 |
| TGME49_310010 | roptry neck protein RON1 | 13 |
| TGME49_215775 | roptry protein ROP8 | 12 |
| TGME49_286420 | elongation factor 1-alpha (EF-1-ALPHA), putative | 9 |
| TGME49_257680 | myosin light chain MLC1 | 8 |
| TGME49_231630 | alveolin domain containing intermediate filament IMC4 | 8 |
| TGME49_316400 | alpha tubulin TUBA1 | 7 |
| TGME49_219320 | gliding-associated protein GAP50 | 7 |
| TGME49_258580 | roptry protein ROP17 | 5 |
| TGME49_319560 | microneme protein MIC3 | 5 |
| TGME49_258870 | cyst wall protein CST7 | 5 |
| TGME49_209910 | histone H2Bv | 4 |
| TGME49_289750 | ribosomal-ubiquitin protein RPL40 | 4 |
| TGME49_308090 | roptry protein ROP5 | 4 |
| TGME49_311240 | DnaJ family chaperone J1 | 4 |
| TGME49_293590 | 3-oxoacyl-acyl-carrier protein synthase I/II, putative | 3 |
| TGME49_203310 | dense granule protein GRA7 | 3 |
| TGME49_271050 | SAG-related sequence SRS34A | 3 |
| TGME49_218260 | histone H3.3 | 3 |
| TGME49_207040 | hypothetical protein | 3 |
| TGME49_235402 | CorA family Mg2+ transporter protein | 3 |
| TGME49_309885 | hypothetical protein | 2 |
| TGME49_313140 | isocitrate dehydrogenase 2 | 2 |

