## Supplementary Figures for "Molecular features of Myosin F adapted for driving actin flows in *Toxoplasma gondii*"

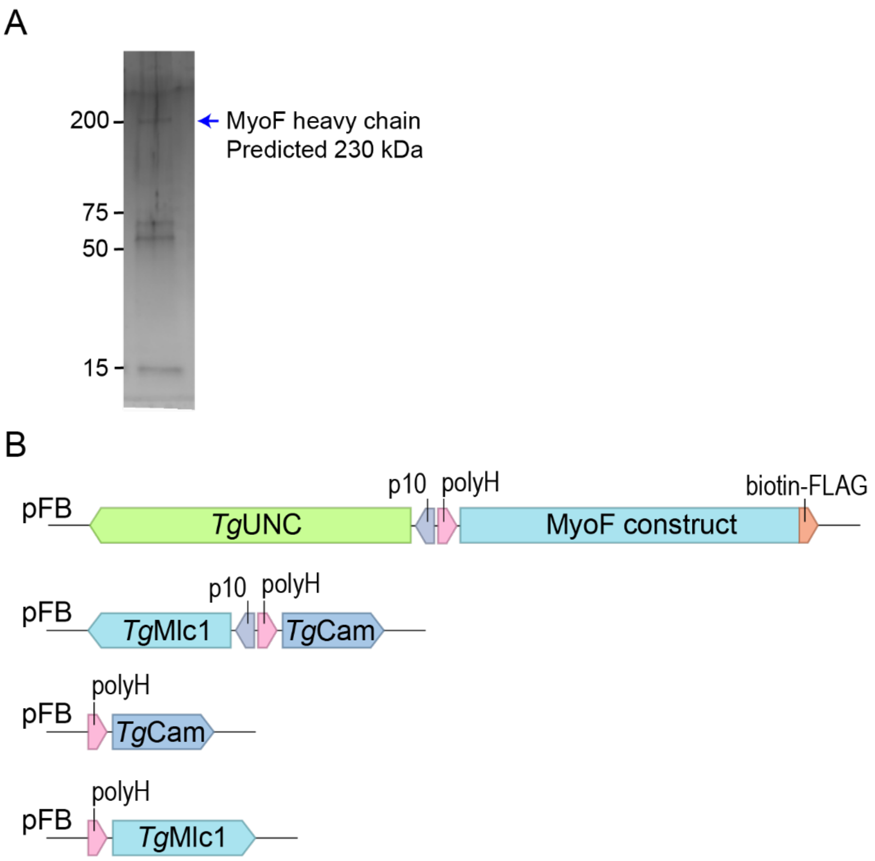


**Figure S1: MyoF immunoprecipitation from *T. gondii* cell extracts and strategy for expressing MyoF constructs in the *Sf9*/baculovirus system.**

(**A**) Representative silver stained SDS-PAGE gel showing the elution following a MyoF-GFP immunoprecipitation assay from *T. gondii* cell lysates using GFP trap affinity resin. (**B**) Schematic of *Sf9* baculovirus constructs for the expression and purification of MyoF constructs, containing a C-terminal biotin-FLAG tag. MyoF constructs were expressed back-to-back with the *T. gondii* chaperone, *Tg*UNC from polyhedrin and p10 promoters. MyoF constructs were co-infected with *Tg*Cam and *Tg*Mlc1 expressed back-to-back or separately depending on the experiment.

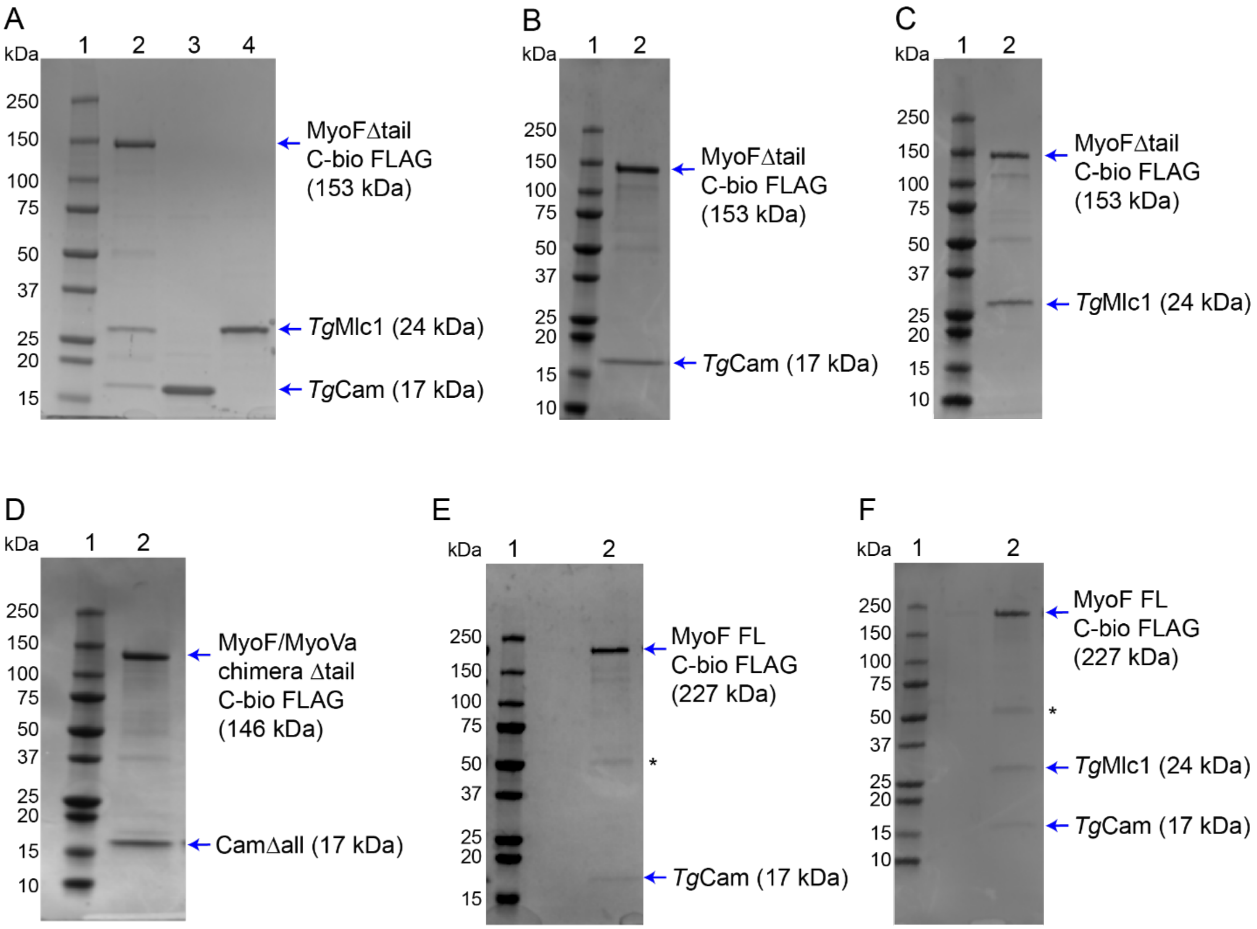


**Figure S2: Coomassie stained SDS-PAGE gels of proteins used in this study.**

(**A**) Protein molecular weight marker (lane 1), MyoFΔtail co-purified with *Tg*Cam and *Tg*Mlc1 from *Sf9* cells (lane 2), HIS purified bacterial expressed *Tg*Cam1 (lane 3) and *Tg*Mlc1 (lane 4). (**B**) Protein molecular weight marker (lane 1), MyoFΔtail co-purified with *Tg*Cam (lane 2). (**C**) Protein molecular weight marker (lane 1), MyoFΔtail co-purified with *Tg*Mlc1 (lane 2). (**D**) MyoF/Va chimeric construct co-purified with mammalian CamΔall (lane 2). (**E**) Protein molecular weight marker (lane 1), full-length MyoF co-purified with *Tg*Cam (lane 2). (**F**) Protein molecular weight marker (lane 1), full-length MyoF co-purified with *Tg*Cam and *Tg*Mlc1. Asterisks indicate contaminating *Sf9* tubulin identified using liquid chromatography-mass spectrometry (LC-MS).

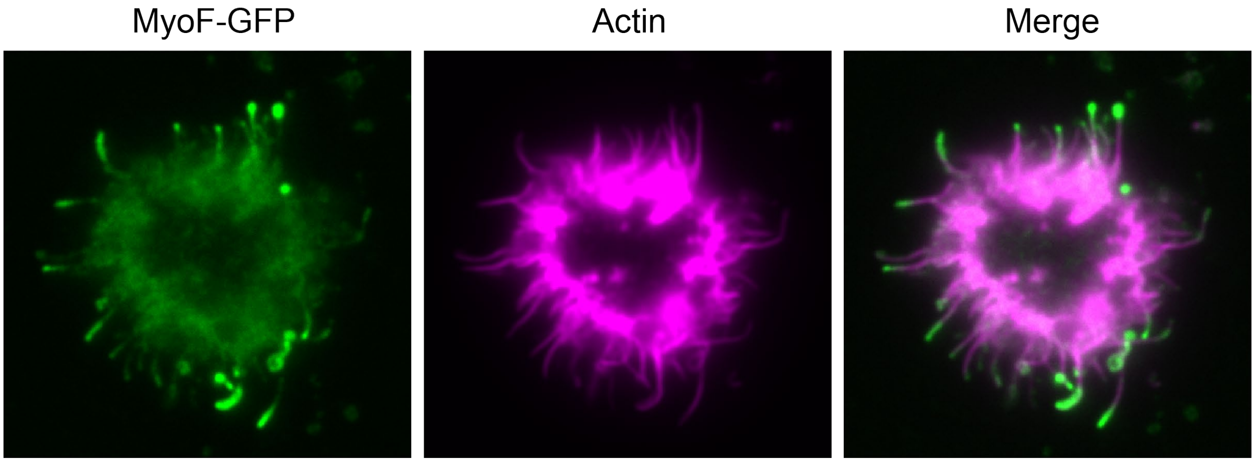


**Figure S3: Ectopic expression of MyoF-GFP in *Sf9* cells.**

Epifluorescence microscopy images showing the localization of full-length MyoF (green) in *Sf9* cells expressing MyoF fused at its C-terminus with an mClover3 variant of GFP. Actin (magenta) is labeled with rhodamine-phalloidin.

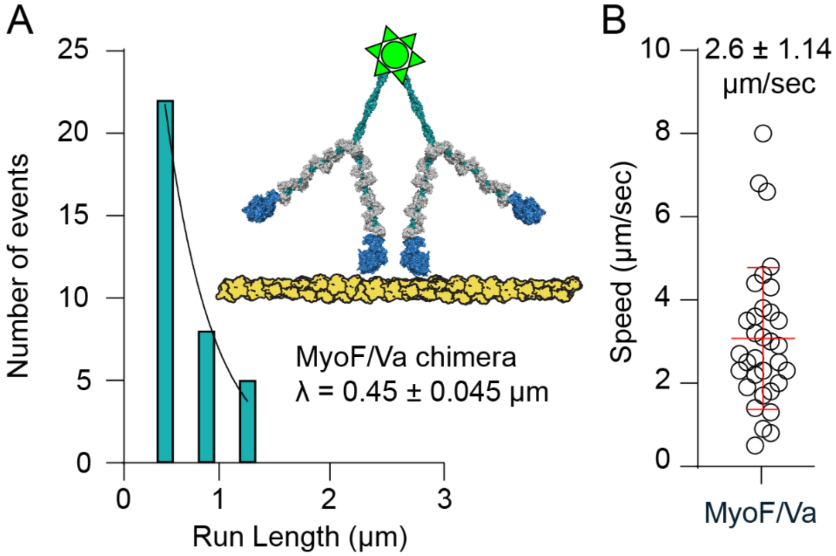


**Figure S4: Motility of small ensembles of MyoF/Va on skeletal vs. *Tg*Act1 actin filaments.**

(**A**) Run length histogram of motor ensembles of the MyoF/Va chimeric construct bound to CamΔall linked through a Qdot on *T. gondii* actin (n = 33). (**E**) Speed distribution of MyoF/Va multiple motor motility on *T. gondii* actin. The mean speed is indicated with a red line. Error is in SD; n = 33.

### Supplementary Movies

**Movie S1.** Gliding filament *in vitro* motility of MyoFΔtail co-purified with *Tg*Cam and *Tg*Mlc1. Frames acquired at 1 sec intervals, 37°C. 15x playback, Image width 32.5 µm.

**Movie S2.** Gliding filament *in vitro* motility of MyoFΔtail co-purified with *Tg*Cam. Frames acquired at 1 sec intervals, 37°C. 15x playback, Image width 32.5 µm.

**Movie S3.** Gliding filament *in vitro* motility of MyoFΔtail co-purified with *Tg*Mlc1. Frames acquired at 1 sec intervals, 37°C. 15x playback, Image width 32.5 µm.

**Movie S4.** Single molecule motility of MyoFΔtail co-purified with *Tg*Cam and *Tg*Mlc1 bound to a streptavidin conjugated Qdot 655 (green) on skeletal actin stabilized with Alexa 488 phalloidin (magenta) (no events observed). Frames acquired at 300 ms intervals, 37°C. 9x playback, Image width 32.5 µm.

**Movie S5.** Movement of multiple motors of MyoFΔtail co-purified with *Tg*Cam and *Tg*Mlc1 bound to a streptavidin conjugated Qdot 655 (green) on skeletal actin filaments (magenta). Frames acquired at 300 ms intervals, 37°C. 4.5x playback, Image width 32.5 µm.

**Movie S6.** Single molecule motility of MyoFΔtail co-purified with *Tg*Cam bound to a streptavidin conjugated Qdot 655 (green) on skeletal actin stabilized with Alexa 488 phalloidin (magenta) (no events observed). Frames acquired at 200 ms intervals, 37°C. 6x playback, Image width 66.6 µm.

**Movie S7.** Movement of multiple motors of MyoFΔtail co-purified with *Tg*Cam bound to a streptavidin conjugated Qdot 655 (green) on skeletal actin stabilized with Alexa 488 phalloidin (magenta). Frames acquired at 50 ms intervals, 37°C. 1x playback, Image width 32.5 µm.

**Movie S8.** Single molecule motility of MyoFΔtail co-purified with *Tg*Cam bound to a streptavidin conjugated Qdot 655 (green) on skeletal actin stabilized with Alexa 488 phalloidin bundled with the addition of a 1:1.1 molar ratio of fascin (magenta) (no events observed). Images were captured using TIRF microscopy. Frames acquired at 50 ms intervals, 23°C. 1x playback, Image width 55 µm.

**Movie S9.** Single molecule motility of full-length MyoF co-purified with *Tg*Cam bound to a streptavidin conjugated Qdot 655 (green) on skeletal actin stabilized with Alexa 488 phalloidin (no events observed). Frames acquired at 200 ms intervals, 37°C. 3x playback, Image width 32.5 µm.

**Movie S10.** Single molecule motility of full-length MyoF co-purified with *Tg*Cam bound to a streptavidin conjugated Qdot 655 (green) on *Tg*Act1 actin filaments stabilized with jasplakinolide and visualized with chromobody fused to EmeraldFP (magenta). Frames acquired at 200 ms intervals, 37°C. 2x playback, Image width 25.1 μm.

**Movie S11.** Single molecule motility of full-length MyoF co-purified with *Tg*Cam and *Tg*Mlc1 bound to a streptavidin conjugated Qdot 655 (green) on *Tg*Act1 actin filaments stabilized with jasplakinolide and visualized with chromobody fused to EmeraldFP (magenta). Frames acquired at 200 ms intervals, 37°C. 3x playback, Image width 32.5 μm.

**Movie S12.** Movement of multiple motors of MyoF/Va chimera co-purified with mammalian CamΔall bound to a streptavidin conjugated Qdot 655 (green) on skeletal actin stabilized with Alexa 488 phalloidin (magenta). Frames acquired at 50 ms intervals, 37°C. 1x playback, Image width 32.5 μm.

**Movie S13.** Movement of multiple motors of MyoF/Va chimera co-purified with mammalian CamΔall bound to a streptavidin conjugated Qdot 655 (green) on *Tg*Act1 filaments stabilized with jasplakinolide and imaged with chromobody fused to EmeraldFP (magenta). Frames acquired at 50 ms intervals, 37°C. 3x playback, Image width 13 μm.

**Movie S14.** Single molecule motility of MyoF/Va chimera co-purified with mammalian CamΔall bound to a streptavidin conjugated Qdot 655 (green) on *Tg*Act1 filaments stabilized with jasplakinolide and imaged with chromobody fused to EmeraldFP (magenta). Frames acquired at 50 ms intervals, 37°C. 3x playback, Image width 32.5 μm.

**Movie S15.** Single molecules of full-length MyoF labeled with Alexa Fluor 647 streptavidin (green) statically associate with Cy5-labeled taxol stabilized microtubules (magenta). Frames acquired at 2 s intervals, 37°C. 16x playback, Image width 32.5 μm.

**Movie S16.** Single molecules of MyoF cc + WD40 labeled with Alexa Fluor 647 streptavidin (green) statically associate with Cy5-labeled taxol stabilized microtubules (magenta). Frames acquired at 2 s intervals, 37°C. 16x playback, Image width 32.5 μm.

**Movie S17.** Dynamic actin-microtubule crosslinking assay showing the relative gliding of Alexa Fluor 488 phalloidin stabilized skeletal F-actin (green) by full-length MyoF (unlabeled) that is bound to a surface immobilized Cy5-labeled microtubule (magenta). Frames acquired at 2 s intervals, 37°C. 40x playback, Image width 36 μm.
